# Too much too many: comparative analysis of morabine grasshopper genomes reveals highly abundant transposable elements and rapidly proliferating satellite DNA repeats

**DOI:** 10.1101/2020.08.22.247130

**Authors:** Octavio M. Palacios-Gimenez, Julia Koelman, Marc Palmada Flores, Tessa M. Bradford, Karl K. Jones, Steven J. B. Cooper, Takeshi Kawakami, Alexander Suh

## Abstract

**Background:** The repeatome, the collection of repetitive DNA sequences represented by transposable elements (TEs) and tandemly repeated satellite DNA (satDNAs), is found in high proportion in organisms across the tree of life. Grasshoppers have large genomes (average 9 Gb), containing large amounts of repetitive DNA which has hampered progress in assembling reference genomes. Here we combined linked-read genomics with transcriptomics to assemble, characterize, and compare the structure of the repeatome and its contribution to genome evolution, in four chromosomal races of the morabine grasshopper *Vandiemenella viatica* species complex.

**Results:** We obtained linked-read genome assemblies of 2.73-3.27 Gb from estimated genome sizes of 4.26-5.07 Gb DNA per haploid genome of the four chromosomal races of *V. viatica*. These constitute the third largest insect genomes assembled so far (the largest being two locust grasshoppers). Combining complementary annotation tools and manual curation, we found a large diversity of TEs and satDNAs constituting 66 to 75 % per genome assembly. A comparison of sequence divergence within the TE classes revealed massive accumulation of recent TEs in all four races (314-463 Mb per assembly), indicating that their large genome size is likely due to similar rates of TE accumulation across the four races. Transcriptome sequencing showed more biased TE expression in reproductive tissues than somatic tissues, implying permissive transcription in gametogenesis. Out of 129 satDNA families, 102 satDNA families were shared among the four chromosomal races, which likely represent a repertoire of satDNA families in the ancestor of the *V. viatica* chromosomal races. Notably, 50 of these shared satDNA families underwent differential proliferation since the recent diversification of the *V. viatica* species complex.

**Conclusion:** In-depth annotation of the repeatome in morabine grasshoppers provided new insights into the genome evolution of Orthoptera. Our TEs analysis revealed a massive recent accumulation of TEs equivalent to the size of entire *Drosophila* genomes, which likely explains the large genome sizes in grasshoppers. Although the TE and satDNA repertoires were rather similar between races, the patterns of TE expression and satDNA proliferation suggest rapid evolution of grasshopper genomes on recent timescales.

## Background

Eukaryotic genomes exhibit repetitive DNA sequences represented by interspersed transposable elements (TEs) and tandem repeats (TRs; e.g., satellite DNA; satDNA), collectively known as the ‘repeatome’ [1]. TEs occupy a large fraction of genomes in organisms throughout the tree of life [2]. The ubiquity of TEs is driven by the ability to either copy and paste themselves with an RNA intermediate step (retrotransposons, class I) or to cut and paste themselves (most DNA transposons, class II) within the genome of the host organism [3–5]. Each TE class can be further categorized by elements that encode protein products required for transposition (autonomous) and those that only contain the sequences (non-autonomous) necessary for trans-recognition by the transposition machinery of an autonomous counterpart [6]. Class I elements comprise SINE, LINE and LTR. Class II elements comprise TIR (DNA transposons), Crypton, RC (rolling-circle replication) and Maverick [7]. The mechanism of transposition allows TEs to invade the genome in a parasitic way without general advantage to the individual carrying them [3] but with potentially deleterious effects on their host by promoting ectopic recombination, mediating chromosomal rearrangements, and disrupting coding sequences [8–10].

Another type of repetitive element widely distributed in eukaryotic genomes is satDNA. It consists of non-coding repetitive DNA that is tandemly arranged and largely represented in the centromeric and pericentromeric heterochromatin of most eukaryotic genomes [11–13]. satDNA evolution is influenced by several mechanisms of non-reciprocal genetic exchange such as unequal crossing-over, intra-strand homologous recombination, gene conversion, rolling-circle replication and transposition [11,14–17]. These mechanisms can gradually increase the copy number of a new sequence variant within a satDNA family across the genomes of a sexual population [11,14–16,18,19]. Sequences within a satDNA family undergo concerted evolution as repeat exchanges occur both within and between members of the satDNA family by non-reciprocal genetic transfers between homologous and sometimes non-homologous chromosomes [14,20]. This results in frequent homogenization of repeats between copies within species and also between repeat copies located on a same chromosome than between different chromosomes [14,20]. At the same time, the primary sequence of satDNAs usually mutates quickly, and this rapid satDNA turnover leads to distinct composition and genomic distribution of satDNAs between strains, populations, subspecies or species [11,15,18,19,21,22]. The library hypothesis proposes that species do not entirely lose or gain certain lineages of satDNAs, but, instead, related species share a common collection of satDNAs that may independently increase or decrease in their copy numbers during or after speciation [23,24]. Sequence divergence as the outcome of reproductive isolation might then lead to a formation of species-specific profiles of satDNA sequence variants [25,26]. For grasshoppers and crickets (Orthoptera), high-throughput sequencing analyses detected 316 satDNA families in ~20 species [12,21,27–32]. Because of the large divergence time between these Orthopteran species (73-224 millions years, Myr) [33], none of these 316 satDNA families showed interspecific homology, except for partial sequence homology within satDNAs of *Gryllus* cricket species [27] (unknown divergence time) and within *Schistocerca* grasshopper species [21] with divergence time < 8 Myr [34]. Because homology between satDNAs often cannot be detected between distantly related species due to rapid sequence divergence, an explicit examination of the library hypothesis requires well-annotated genomes of closely related species.

The repeatome is involved in the processes of sex chromosome differentiation [11,35–38]. Both plants and animals have accumulated TEs and satDNAs in the non-recombining regions of Y and W chromosomes [11,35,37–41]. Studies on neo-Y chromosomes of several *Drosophila* species proposed that the first steps of Y chromosome degeneration are driven by accumulation of TEs and satDNAs [36,40]. The non-recombining parts of the Y or W chromosomes may thus expand by repeat accumulation and heterochromatinization [36,40]. Moreover, transcription of satDNAs (satRNAs) may have a critical role in centromere function, chromatin silencing, heterochromatin formation, chromatin modulation, and up-regulation of X-linked genes in chromosome dosage compensation [42–45]. It has been proposed that interspecific incompatibilities in hybrids between satRNAs and specific proteins involved in centromere function can contribute to genome divergence and the speciation process [25,44,46–48].

Grasshoppers generally have large genomes (9 Gb on average, minimum 1.5 Gb and maximum 16.6 Gb [49]) likely because of large amounts of repetitive DNA [50,51]. They provide ample opportunities to investigate the influence of the repeatome on karyotype evolution because grasshoppers are also karyotypically divergent between closely related species [52–54]. However, comparative genomic studies in grasshoppers have been hampered by their large genome sizes [50,51], The Australian morabine (Morabinae) grasshopper of the genus *Vandiemenella* (hereafter referred to as the *viatica* species group) is a relatively young species complex with estimated divergence time < 0.5-3.1 Myr based on a mitochondrial marker [53] and is karyotypically diverse [53–55]. It currently contains two nominal species (*V. pichirichi* and *V. viatica*) and five provisional species (P24, P25, P45b, P45c, and P50) differentiated by one or more chromosomal rearrangements [56,57], Earlier cytogenetics studies hypothesized that viatica19 (2n = 19, X0 male) is most closely resembling the ancestral karyotype of the *viatica* species group [54,55]. Subsequent sequential chromosomal rearrangements, including centric fusions, fissions and inversions, resulted in the formation of the present taxa [54,55,58], Notably, neo-sex chromosomes in the *viatica* species group evolved three times independently through recent fusions of the ancestral X chromosome with a different autosome each time (P24X0/XY, P25X0/XY and P45bX0/XY races) [54,55], Given that repetitive sequences can play critical roles in the evolution of genome structure and function, a comprehensive analysis of TEs and satDNAs of these grasshopper genomes is essential for understanding of the genome structure and chromosomal evolution of the *viatica* species group.

Here, we characterized TEs and satDNAs in four chromosomal races of the *viatica* species group, P24X0, P24XY, P45bX0, and P45bXY, by generating 10X Genomics Chromium linked-read data and RNA sequencing data. We then used three complementary methods, namely homologybased, structure-based, and de-novo approaches, for annotating the TE and satDNA fraction of the genomic reads and assemblies. By comparing sequence divergence within the TE classes, we identified the temporal dynamics of TE accumulation. We identified transcriptional activity of many subfamilies of TEs in three tissues of these grasshoppers, showing that the TE expression levels vary greatly among TEs, tissues and sexes. We also provided evidences that satDNAs expanded and contracted in their genomic copy numbers at different time points since the divergence of the chromosomal races.

## Results

### Genome assembly

We sequenced male genomes of four chromosomal races representing two pairs of karyotypes with and without neo-sex chromosomes (Figure 1a) using 10X Genomics Chromium linked-read libraries with 1,577-1,883 million paired-end reads per library. Average input molecule lengths between races ranged from 17.44 to 56.11 kb. By using Supernova 2.1.0 [59], we obtained genome assemblies with the following sizes: 3.02 Gb in P24X0, 2.73 Gb in P24XY, 3.27 Gb in P45bX0, and 2.94 Gb in P45bXY. The contig N50 for genome assemblies ranged from 29.11 to 35.69 kb, and the scaffold N50 ranged from 34.85 to 316.69 kb. Chromosomal races with larger inferred genome assembly sizes tended to have more fragmented genomes (Table 1). Using BUSCO v3 [60] with the Arthropoda dataset, these assemblies had more complete single-copy orthologs (57-90 %) than the Illumina assembly of the migratory locust *Locusta migratoria* (Acrididae, Oedipodinae) (42 %) [50]. The BUSCO scores also indicated that the fractions of genes that were entirely missing in each race (except P45bXY) were lower than what has been observed previously in *L. migratoria* genome assembly (Figure 1b). The assembly of P45bXY showed the lowest number of complete single copy orthologs across the *V. viatica* genome assemblies but still higher than the *L. migratoria* genome assembly [50].

**Figure 1.**
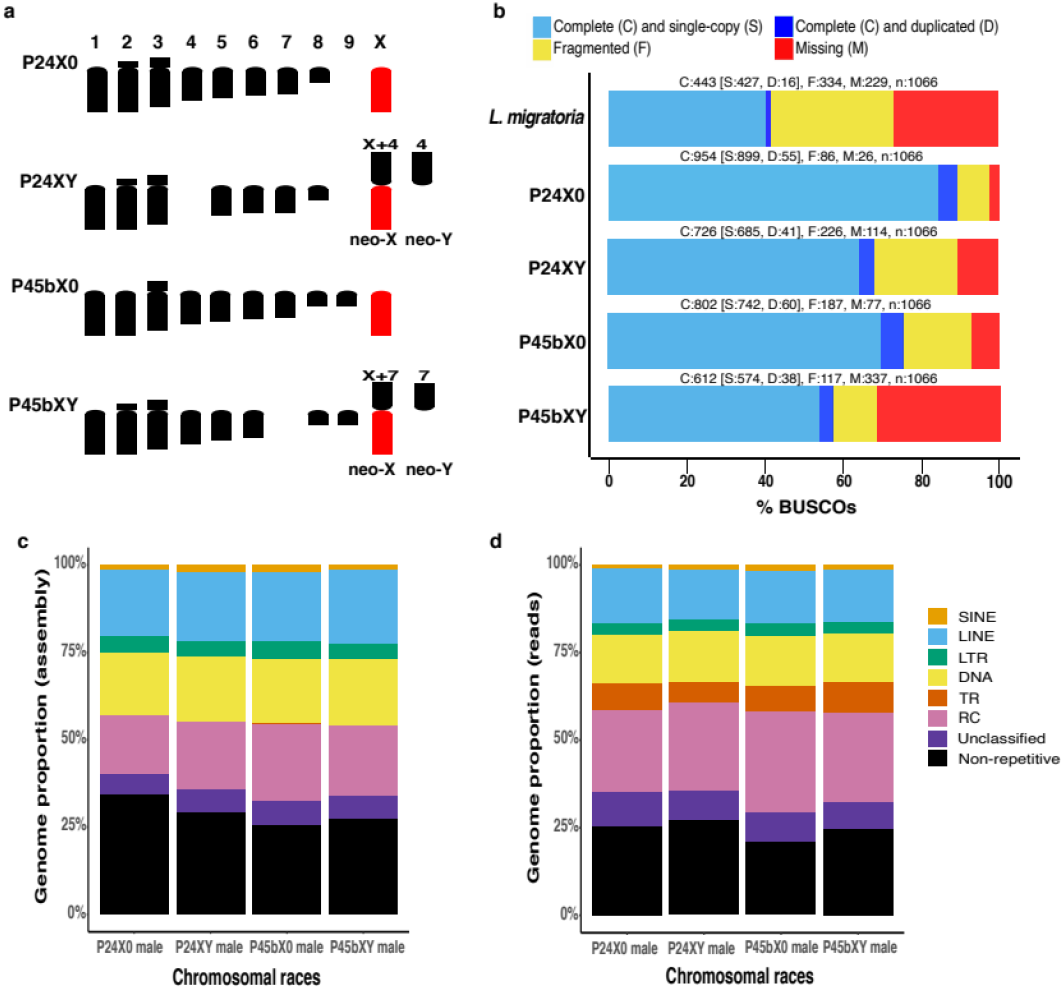
Overview of the karyotype, BUSCO assembly assessment, and repeat composition across four chromosomal races of the *viatica* species group. **a)** Karyotypes of the four chromosomal races. Chromosomes are aligned based on centromere position. Only haploid sets of chromosomes in males are shown. Two independent emergences of neo-XY sex chromosomes via fusions between the ancestral X chromosome (red) and one of the autosomes (black) are highlighted. **b)** BUSCO completeness assessment for genome assembly quality control. Color-coded bar plot showing the proportions of BUSCO genes classified as complete, complete single-copy, complete duplicated, fragmented, and missing. **c-d)** Color-coded bar plot illustrating the proportions of major TE groups, unclassified repeats, tandem repeats (TRs), and non-repetitive regions detected in the assembled genomes **(d)** and in the sequenced reads **(d)**. The major TE groups included manually curated RepeatModeler consensus sequences and Helitrons (RC, rolling circle replication) identified by HelitronScanner.

**Table 1:** Pseudohaploid assembly statistics across the four chromosomal races of the *viatica* species group.

|  | **P24X0** | **P24XY** | **P45bX0** | **P45bXY** |  |
| --- | --- | --- | --- | --- | --- |
|  | **male** | **male** | **male** | **male** |  |
| Total millions of reads (2x150 bp) | 1,883 | 1,582. | 1,759 | 1,577 | 10X Genomics Chromium linked reads |
| Molecule length (kb) | 41.18 | 17.44 | 56.11 | 30.44 | Average input molecule lengths |
| Long scaffold (kb) | 44.43 | 100.68 | 88.04 | 99.29 | Number of scaffolds >= 10 kb |
| Scaffold N50 (kb) | 316.69 | 34.85 | 51.83 | 36.43 | N50 scaffold size |
| Edge N50 (kb) | 8.44 | 6.24 | 8.24 | 6.73 | N50 edge size |
| Contig N50 (kb) | 35.69 | 26.94 | 33.34 | 29.11 | N50 contig size |
| Phaseblock N50 (kb) | 651.54 | 50.13 | 115.56 | 53.56 | N50 phase block size |
| Assembly GC content % | 37.99 | 37.93 | 38.09 | 38.01 |  |
| Assembled genome size (Gb) | 3.02 | 2.73 | 3.27 | 2.94 | Only scaffolds >= 10 kb |
| Genome size estimation (Gb) | 3.30 | 3.36 | 3.52 | 3.82 | Supernova (computed from k-mer distributions) |
|  | 5.42 | 5.65 | 5.76 | 6.32 | FindGSE (computed from k-mer distributions) |
| **Average genome size (Gb)** | **4.26** | **4.50** | **4.64** | **5.07** | Based on k-mer distributions |

Owing to the incompleteness of virtually all animal genome assemblies [61,62], genome assembly length tends to be smaller than the actual genome size. To ameliorate this underestimation, we inferred genome size from sequencing reads directly by analyzing the frequency of k-mers using the findGSE function [63]. The measures of genome sizes varied between each pipeline used, i.e. between findGSE and the Supernova estimations computed from k-mer distributions (Table 1). We thus report average values between these k-mer-based measures as an approximation of the true genome sizes. This analysis resulted in genome size estimates of 4.26 Gb in P24X0, 4.50 Gb in P24XY, 4.64 Gb in P45bX0, and 5.07 Gb in P45bXY.

### Transposable element (TE) identification

We first ran RepeatModeler 1.0.8 [64] on each of the four genome assemblies to generate benchmark repeat libraries for annotating the TEs, yielding between 1,361 and 1,398 consensus sequences per race. Between 637 and 668 consensus sequences were initially classified as unknown by RepeatModeler in each library. To further classify these unknown repeats, we manually curated the P24X0 repeat library generated by RepeatModeler because this had the best genome assembly quality based on contig N50 and BUSCO scores. Our manual curation identified 212 new consensus sequences of TE subfamilies in the P24X0 genome (32 % of the unknown repeat consensus sequences above). Next, we used the curated TE consensus sequences of P24X0 to re-classify unknown repeats in the other three libraries by homology searches in RepeatMasker 4.0.8 [65]. This classified a total of 215, 206, and 212 new TE subfamilies (~34 % of the unknown repeat consensus sequences above) in the P24XY, P45bX0, and P45bXY genomes, respectively. From these newly identified TEs, about 2.24-2.69 % of each genome assembly (83-93 subfamilies) were reclassified as LTR retrotransposons and 2.13-2.78 % of each genome assembly (77-98 subfamilies) as DNA transposons. The comparison of TE landscapes, i.e., the distribution of TE-derived bp in bins of Kimura 2-parameter (K2P) distance, in raw RepeatModeler repeat libraries vs. curated libraries highlights the improvement of TE annotation by in-depth manual curation and re-classification (Additional file 2: Figure S1).

To further improve the TE annotation, we applied HelitronScanner 1.0 [66] to search for low-copy Helitrons that were missed by RepeatModeler. HelitronScanner and clustering of the nucleotide dataset into clusters that met a similarity threshold of 80 % produced a set of non-redundant representative sequences (families) of new Helitrons, i.e. 230, 211, 286, and 221 Helitron families in the P24X0, P24XY, P45bX0, and P45bXY genomes, respectively. The bp percentages of these new Helitron sequences in the genome assemblies were 10 % (2,743,812 fragments) in P24X0, 13 % (2,578,114 fragments) in P24XY, 15 % (3,207,327 fragments) in P45bX0, and 13 % (2,837,154 fragments) in P45bXY. We identified a new family of autonomous Helitron (“Tukutron”) in the P24X0 genome (Figure 2) in which entire and truncated SINEs were nested in its sequence. Tukutron comprises 12,053 fragments in the genome of P24X0 (0.01 % of the genome).

**Figure 2.**
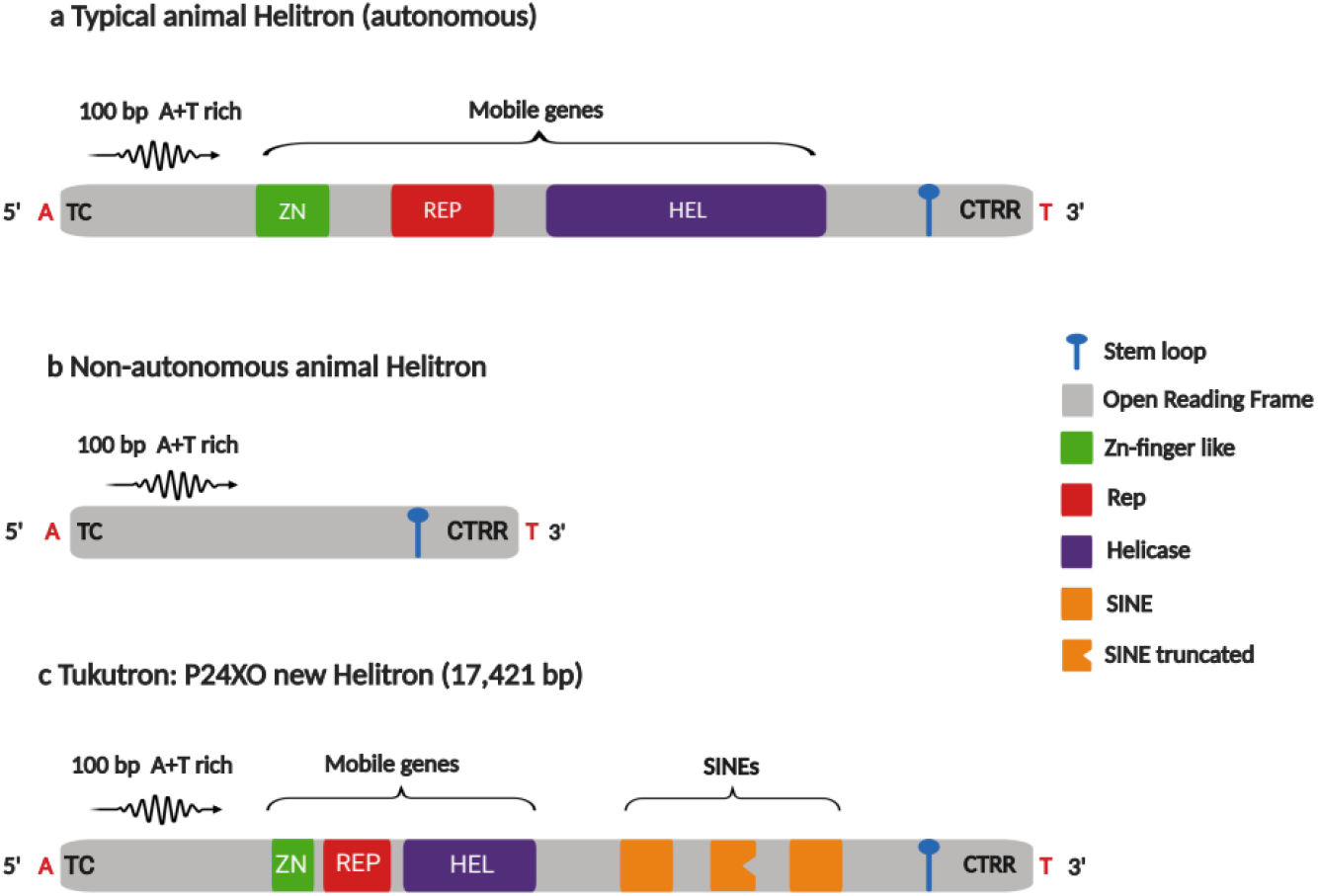
Helitron structures found in the genome assemblies of the *viatica* species group. **a)** Typical autonomous animal Helitron. **b)** Non-autonomous animal Helitron. **(c)** The new identified Helitron family identified in the P24X0 chromosomal race, named Tukutron. The new Helitron bears entire and truncated copies of short interspersed elements (SINEs) which are similar to SINEs in the *Locusta migratoria* genome (SINE2-3_Lmi in Repbase).

The above TE annotations revealed that from 66 to 75 % of the assembled genomes of the *viatica* species group was composed of TEs (Figure 1c, Table 2). The density of LINEs (715 to 819 Mb) was highest among the annotated major TE groups, followed by rolling-circle (RC) Helitrons (664 to 878 Mb), DNA transposons (679 to 734 Mb), LTRs (160 to 208 Mb) and SINEs (56 to 87 Mb) (Table 2). Regarding superfamilies of TEs, the most abundant across the assembled genomes was RC/Helitron (17-22 %), followed by DNA/TcMar (7%-9%) and LINE/CR1 (7-10 %). Since tandem repeats (TRs) were likely underrepresented in genome assemblies, the read-based RepeatExplorer2 [67–69] and NOVOplasty 3.7.2 protocols [70] were applied to detect TRs (see below). None of the TRs detected using these two approaches were recovered by the above RepeatModeler repeat libraries. The relative genomic abundance of detected TRs in the sequenced reads was then compared by sampling 4 million read pairs per library and aligning them to the combined repeat libraries database (RepeatModeler + HelitronScanner + TRs) with RepeatMasker; this quantification was done separately in each race. The analysis revealed that 73 to 79 % of the sequenced reads of the *viatica* species group was composed of repeats (Figure 1d, Table 3), and showed that, while overall TE proportions were comparable to the assembly proportions, TRs and RC/Helitron sequences were better represented in the read-based approach than in the assembly-based approach (Figure 1c,d; Table 2 and 3).

**Table 2.** Assembly-based quantification of repeats. Copy number, total base pair and density of different classes of repeats annotated by RepeatMasker using a combined library of the RepeatModeler *de-novo* library from each race (manual curation of the P24X0 library, used for re-classification of the three other *de-novo* libraries), the Arthropod library from Repbase, and libraries from HelitronScanner and RepeatExplorer2, across males of four chromosomal races of the *viatica* species group. TR = tandem repeats. GP = genome proportion (% assembly).

| Repeat type | P24X0 |  |  | P24XY |  |  | P45bX0 |  |  | P45bXY |  |  |
| --- | --- | --- | --- | --- | --- | --- | --- | --- | --- | --- | --- | --- |
|  | Copies | Total bp | GP | Copies | Total bp | GP | Copies | Total bp | GP | Copies | Total bp | GP |
| SINE | 355,321 | 57,777,060 | 1.45 | 476,672 | 84,209,572 | 2.31 | 498,884 | 86,588,794 | 2.16 | 331,980 | 55,473,057 | 1.45 |
| LINE | 1,630,136 | 751,701,393 | 18.88 | 1,544,315 | 715,290,166 | 19.6 | 1,603,310 | 792,142,166 | 19.79 | 1,780,161 | 819,481,256 | 21.40 |
| LTR | 270,520 | 191,052,819 | 4.80 | 298,156 | 160,264,290 | 4.39 | 316,814 | 207,702,027 | 5.19 | 303,192 | 169,275,629 | 4.42 |
| DNA | 2,269,382 | 712,386,285 | 17.89 | 2,136,964 | 678,964,713 | 18.61 | 2,179,781 | 734,004,927 | 18.34 | 2,165,693 | 714,794,492 | 18.66 |
| RC | 3,194,220 | 663,607,802 | 16.72 | 8,654,890 | 708,282,512 | 19.45 | 3,887,696 | 878,010,926 | 21.88 | 3,569,564 | 782,636,309 | 20.45 |
| TR | 5,269 | 790,157 | 0.02 | 3,294 | 506,934 | 0.01 | 10,988 | 1,403,691 | 0.04 | 232 | 12,409 | 0.01 |
| Unclassified | 889,785 | 237,465,164 | 5.98 | 795,764 | 240,830,299 | 6.62 | 946,725 | 290,976,813 | 7.29 | 8,992,548 | 237,984,790 | 6.24 |
| **Total** | **8,614,633** | **2,614,780,680** | **65.74** | **13,910,055** | **2,588,348,486** | **70.99** | **9,444,198** | **2,989,425,653** | **74.69** | **17,143,370** | **2,779,657,942** | **72.62** |

**Table 3.** Read-based quantification of repeats by sampling 4 million of read pairs per library. Copy number, total base pair and density of different classes of TEs by RepeatMasker using a combined library of the RepeatModeler *de-novo* library from each race (manual curation of the P24X0 library, used for re-classification of the three other de-novo libraries), the Arthropod library from Repbase, and libraries from HelitronScanner and RepeatExplorer2, across males of four chromosomal races of the *viatica* species group. TR = tandem repeats. GP = genome proportion (% reads).

| Repeat type | P24X0 | P24XY | P45bX0 | P45bXY |
| --- | --- | --- | --- | --- |
|  | GP | GP | GP | GP |
| SINE | 1.23 | 1.49 | 1.81 | 1.27 |
| LINE | 15.48 | 14.26 | 14.91 | 15.09 |
| LTR | 3.39 | 3.29 | 3.81 | 3.28 |
| DNA | 13.83 | 14.43 | 13.95 | 13.77 |
| RC | 23.17 | 24.84 | 28.73 | 25.67 |
| TR | 7.82 | 6.06 | 7.53 | 8.85 |
| Unclassified | 9.86 | 8.65 | 8.50 | 7.51 |
| **Total** | **74.78** | **73.02** | **79.44** | **75.44** |

### Temporal accumulation of TEs

Assuming that the K2P distance from the consensus sequence reflects the time since the insertion of a TE copy, this can be a proxy for the temporal accumulation of TEs [71]. Based on this assumption, we quantified the accumulation of recent TEs that were between 0 and 5 % diverged from the respective consensus sequence (Figure 3). We found a total length of 314-463 Mb of TEs with 0-5 % divergence in each assembly, suggesting massive recent amplifications of the five major TE groups in each race: P24X0 (DNA = 175 Mb, LINE = 126 Mb, SINE = 7 Mb, LTR = 42 Mb, RC = 35 Mb), P24XY (DNA = 167 Mb, LINE = 97 Mb, SINE = 2 Mb, LTR = 25 Mb, RC = 23 Mb), P45bX0 (DNA = 200 Mb, LINE = 158 Mb, SINE = 7 Mb, LTR = 56 Mb, RC = 43 Mb) and P45bXY (DNA = 181 Mb, LINE = 122 Mb, SINE = 14 Mb, LTR = 36 Mb, RC = 47 Mb). This is larger than the estimated genome size of *Drosophila* and many other insects [49].

**Figure 3.**
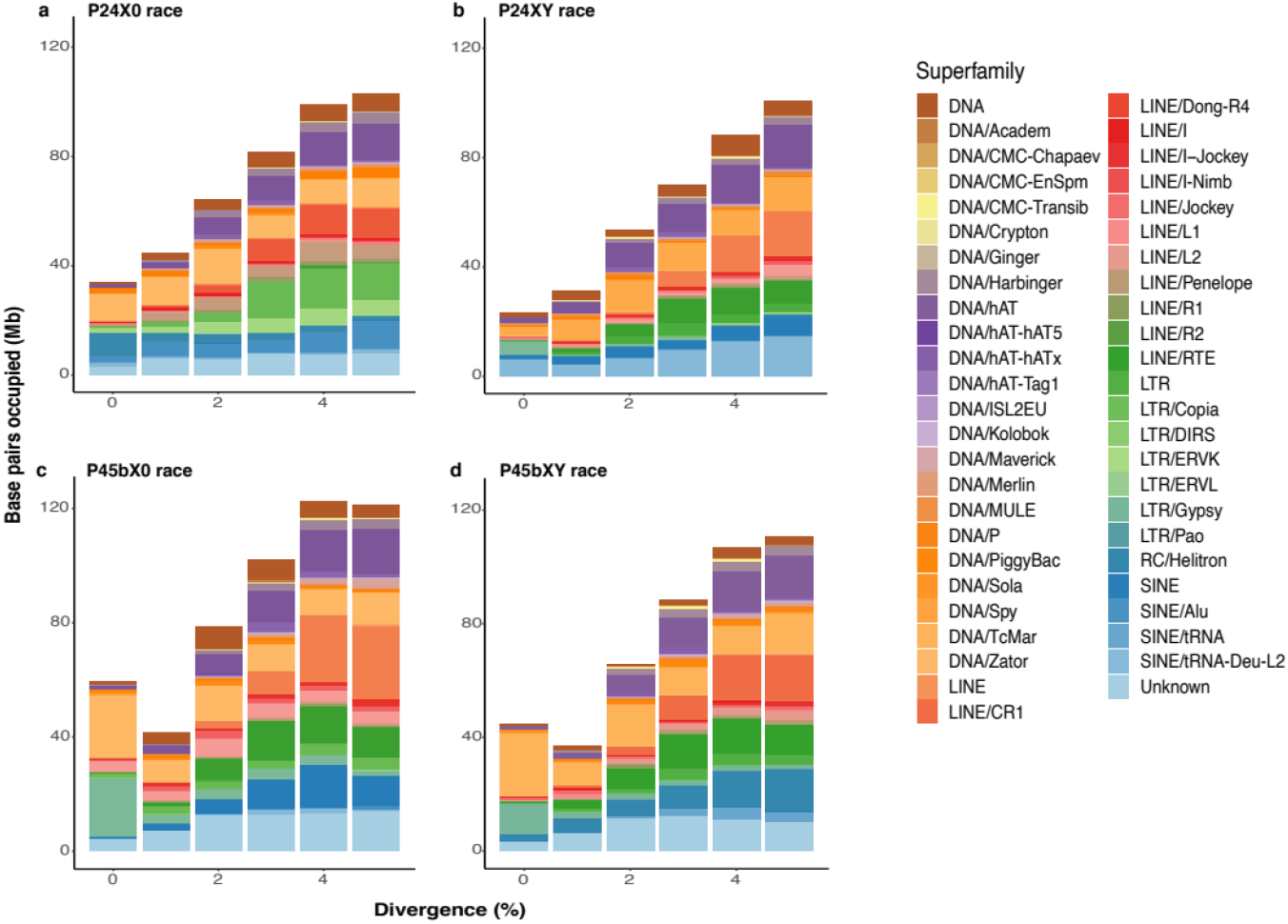
Repeat landscapes illustrating recent accumulation of 49 superfamilies of transposable elements across four chromosomal races of the viatica species group. **a)** P24X0. **b)** P24XY. **c)** P45bX0. **d)** P45bXY. Color-coded bar plots were generated from RepeatMasker .align files after selection of all TE copies with K2P distances < 5 %. The graphs represent the base pairs occupied of a given TE superfamily (*y* axis) in the different genomes analyzed, binned according to K2P distances to their corresponding consensus sequence (*x* axis, K2P distance from 0 to 5%). We considered copies in the divergence bins < 5 % as recent TEs, likely corresponding to copies of recently active elements. Plots showing all divergence bins (K2P distance from 0 to 50 %) are shown in Additional file 2: Figure S1.

We considered the K2P distance bins of 0 to 1 % as very recent TEs that likely accumulated during or after divergence of the chromosomal races. Some of the most abundant TE superfamilies with very recent copies were DNA/DNA, DNA/P, DNA/Sola, DNA/hAT, DNA/TcMar, RC/Helitron, LINE/L2, LTR/LTR, LTR/Gypsy and SINE/tRNA (Figure 3). The numbers of occupied bp of these superfamilies in each assembled genome were different in this divergence bin, with largest variation for LTR/Gypsy (e.g., P24XY = 5 Mb, P24X0 = 9 Mb, P45bXY = 11 Mb, P45bX0 = 21 Mb; coefficient of variation, cv = 170 %) and DNA/TcMar (e.g., P24XY = 3. Mb, P24X0 = 10 Mb, P45bX0 = 22 Mb, P45bXY = 22 Mb; cv = 154 %) indicating differential amplification.

### Transcriptional activities of TEs

We next characterized TE expression in the P24X0 and P24XY races (5-11 individuals per tissue/sex/race) by comparing the number of RNA-seq reads mapping to recent TE copies (i.e., copies with K2P distance < 5 %; hereafter referred to as recent TE expression’ data), using RepEnrich2 (https://github.com/nerettilab/RepEnrich2) and DESeq2 1.20.0 [72]. For comparison, we repeated the analysis with all TE copies regardless of K2P distance. For this analysis, we provided RepEnrich2 with a filtered RepeatMasker annotation file containing only TE loci with < 5 % K2P from the consensus sequences. This step was required to survey recent TE expression because RepEnrich2 does not retain locus coordinates, preventing us to subsample recent TE expression if the total TE expression is considered (i.e., including older TE copies with K2P > 5 %).

In the analysis of recent TE expression, there were 598 and 1,415 expressed TE subfamilies with nonzero total read count in reproductive tissues (male testis and female ovary) in P24X0 and P24XY, respectively. Of these, we observed a higher proportion of female-biased TEs (FBTEs) relative to male-biased TEs (MBTEs) in P24X0 (FBTEs 15.05 %, MBTEs 10.37 %); P24XY showed the opposite trend between sexes (MBTEs 4.88 %; FBTEs 3.39 %) (Table 4). We found 574 and 1,415 expressed TE subfamilies with nonzero total read count in head tissues of P24X0 and P24XY, respectively. Of these, we observed a higher proportion of MBTEs relative to FBTEs in P24X0 (MBTEs 3.66 %, FBTEs 0.35 %) and P24XY (MBTEs 1.48 %, FBTEs 0.21 %) (Table 4). In leg tissues, a total of 1,362 and 775 expressed TE subfamilies with nonzero total read count were observed in P24X0 and P24XY, respectively. There was not much variation in TE expression levels in leg between sexes in both races: P24XY (MBTEs 0 %, FBTEs 0.52 %) and P24X0 (MBTEs 0.23 %, FBTEs 0.07 %). The heatmaps showing the expression data of the 50 most highly expressed TE subfamilies in three tissues contain representatives from all five major TE groups (Figure 4). The numbers of differentially expressed TEs between sexes and tissues when including all TE copies regardless of K2P distance were much larger (between 2,354 and 7,240 TEs transcribed; Additional file 1: Table S1) than the recent TE expression data, likely because of transcription of old and inactive TE copies.

**Figure 4.**
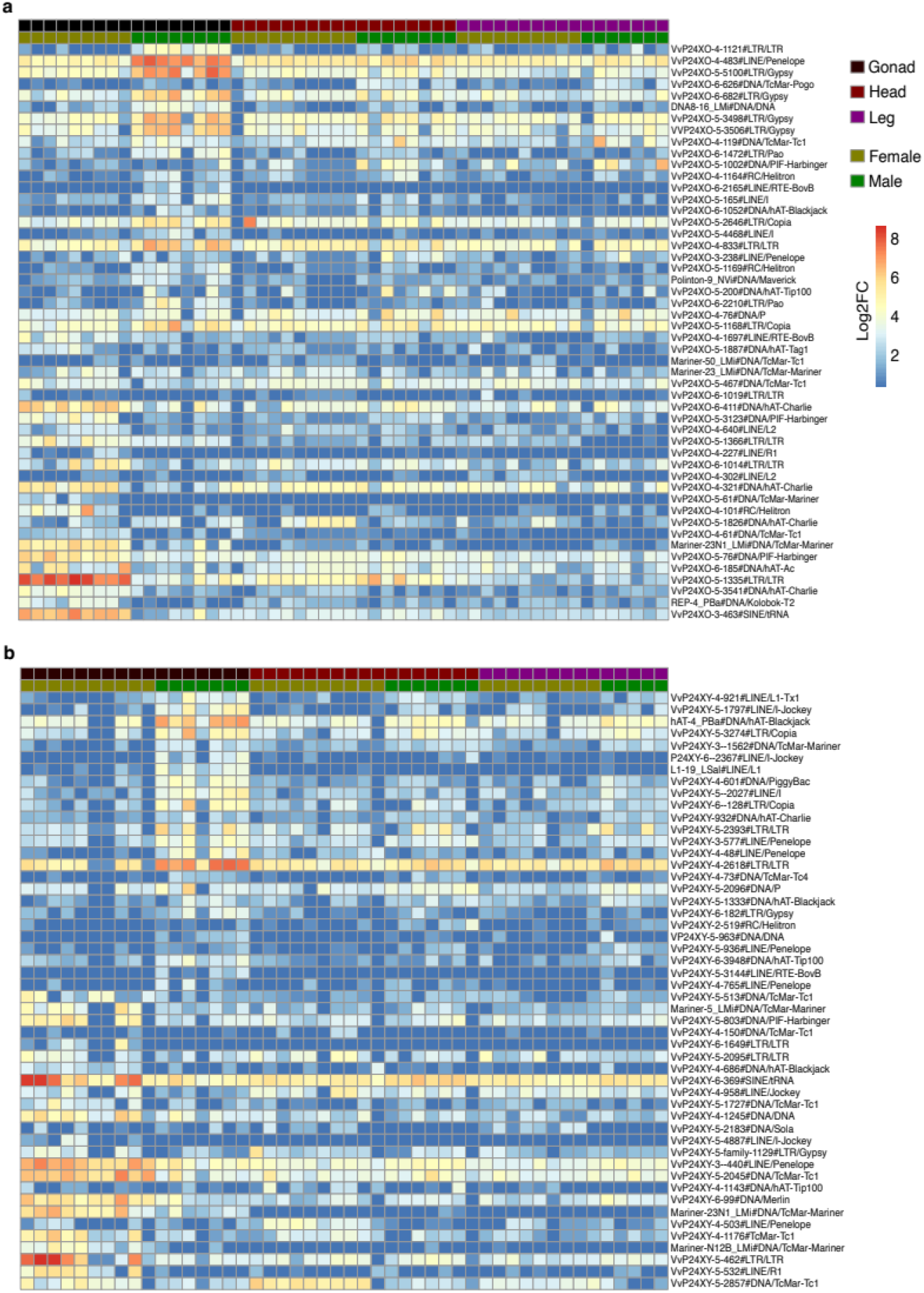
Heatmaps showing RNA-seq expression of the 50 most highly expressed TE subfamilies across sexes and tissues of two chromosomal races of the *viatica* species group. **a)** P24X0. **b)** P24XY. Expression levels shown as log2-normalized counts (log2 fold-change, *P* < 0.05). The color-coded bar indicates the expression levels in each sex and tissue. Each row represents a TE consensus sequence (i.e., subfamily) and each column the biological replicates. The analysis represents recent TE expression based on the TE copies with K2P < 5 % of divergence to their corresponding TE consensus sequence.

**Table 4.** Differentially expressed TE subfamilies containing only recent TEs (genomic copies with < 5 % K2P distance from the consensus sequences) between sexes and tissues in two chromosomal races of the *viatica* species group.

| Race | Sample | No. of<br>expressed TEs | SBTEs (%) | MBTEs (%) | FBTEs (%) |
| --- | --- | --- | --- | --- | --- |
| P24X0 | Head | 574 | 23 (4.05) | 21 (3.66) | 2 (0.35) |
|  | Gonad | 598 | 152 (25.42) | 62 (10.37) | 90 (15.05) |
|  | Leg | 1,362 | 4 (0.29) | 3 (0.23) | 1 (0.07) |
| P24XY | Head | 1,415 | 24 (1.71) | 21 (1.48) | 3 (0.21) |
|  | Gonad | 1,415 | 117 (8.27) | 69 (4.88) | 48 (3.39) |
|  | Leg | 775 | 4 (0.52) | 0 | 4 (0.52) |

We report the total number of TEs expressed, and those TEs that were sex-biased with $\log_2\text{FC} > 0$ and $\log_2\text{FC} < 0$ and adjusted $P < 0.05$ ; $\log_2\text{FC} = \log_2$ fold-change. SBTEs = sex-biased TEs; MBTE = male-biased TEs.

### Tandem repeat (TR) identification and sequence characterization

Since TRs were underrepresented in the aforementioned linked-read assemblies, we applied two read-based approaches, RepeatExplorer2 [67–69] and NOVOplasty 3.7.2 [70], and merged the outputs of these two to detect TRs in each race. We identified 56 TR families in P24X0, 60 TR families in P24XY, 71 TR families in P45bX0 and 92 TR families in P45bXY. These TRs included multigene families (45S rRNA, 5S rRNA, U snRNA, and histones genes), the tandem telomere repeat (TTAGG)_n_, and satDNA families. None of the TRs were found in the aforementioned RepeatModeler libraries. The total abundance of TRs composed of multigene families, telomere repeats and satDNAs represented 7.82 %, 6.06 %, 7.53 % and 8.85 % of the sequenced reads of P24X0, P24XY, P45bX0, and P45bXY, respectively (Table 3, Additional file 1: Table S2-S5). Compared to these read-based TR quantifications, the abundance of TRs in the genome assemblies was much lower (i.e. << 0.1 % per assembly), suggesting that most satDNA copies were collapsed during the assembly process. Read-based approaches thus proved essential to the identification and quantification of TRs (Table 2 and 3).

The telomere repeat was more abundant in the XY races (P24XY and P45bXY, 1.61 % and 1.21 % of the sequenced reads, respectively) than the X0 races (P24X0 and P45bX0, < 0.91 % of the sequenced reads) likely through independent amplification of the telomere repeats in the two former (Figure 5). The 45S rRNA gene underwent slightly more differential amplification in P24XY (abundance 0.61 %) than the other three races (abundances < 0.31 %). Most copies of the telomere repeat and the 45S rRNA gene were in the divergence bins of K2P < 5 %, suggesting recent amplification.

**Figure 5.**
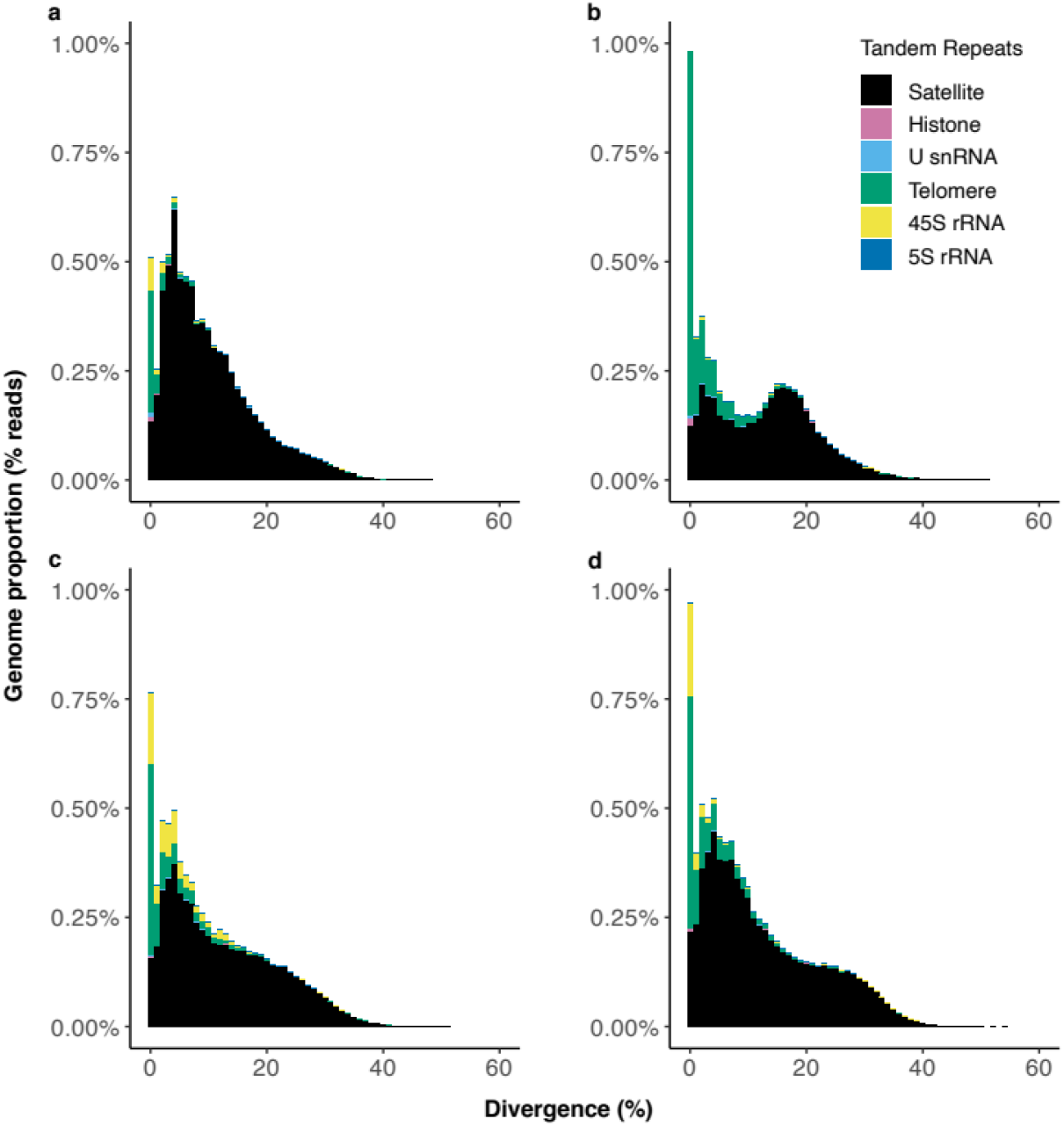
Repeat landscapes of the main groups of tandem repeats (TRs) detected in the sequenced reads of the four chromosomal races of the *viatica* species group. **a)** P24X0. **b)** P24XY **c)** P45bX0. **d)** P45bXY. Color-coded bar plots are based on RepeatMasker showing K2P distance to their corresponding consensus sequence on the *x* axis and the TR abundance on the *y* axis. Color-coded bar plots show the proportions of TRs classified as satellite (45-81 families), telomere repeat and the multigene families U snRNA genes (U1, U2, U5, U6), histone genes (H1, H2A, H2B, H3, H4), 45S rRNA and 5S rRNA genes. More detailed repeat landscapes of all TR families are shown in the Additional file 2: Figure S2.

The satDNAs were the dominant TRs in all four races (Figure 5). RepeatExplorer2 found 45 satDNA families in P24X0 (7 % of sequenced reads), 48 satDNA families in P24XY (4 % of sequenced reads), 60 satDNA families in P45bX0 (6 % of sequenced reads), and 81 satDNA families in P45bXY (7 % of sequenced reads) (Additional file 1: Table S2-S5; Additional file 2: Figure S2). The repeat unit length (monomers) of the satDNA families ranged from 21 to 1,740 bp in P24X0, from 7 to 1,690 bp in P24XY, from 13 to 1,710 bp in P45bX0, and from 13 to 1,250 bp in P45bXY. Multiple sequence alignments and dotplots showed that most satDNA families were present in multiple contigs within the clusters of the RepeatExplorer2 results, with monomers differing by at least one nucleotide from each other, indicating that distinct sequence variants of monomers and higher-order repeat structures (HORs) were present in each genome (Additional file 1: Table S2-S5; Additional file 2: Figure S3).

### Assessment of the satDNA library hypothesis

Because RepeatExplorer2 can only detect satDNAs with abundance > 0.01 % of the genome [67–69], homologous satDNAs present below this threshold can remain undetected across samples. Thus, we used two approaches to search for homologous satDNAs between races. First, we built a satDNA database by concatenating all the satDNA consensus sequences detected in each race (totaling 234 consensus sequences), and performed an all-against-all comparison of the consensus sequences in the database using the rm.homology.py script (https://github.com/fjruizruano/ngs-protocols) [31]. Second, we mapped the genomic reads of each race separately to the satDNA database using RepeatMasker. These approaches defined 129 satDNA families across the four races (hereafter referred to as satDNA-1 to satDNA-129). Out of the 129 satDNAs, 102 satDNAs were shared between all four races, and 27 satDNAs were present or absent in one or more race (Additional file 1: Table S6-S7). From these 27 satDNAs families, there were 1-2 satDNA families specific to just one race each, the largest number in P24XY and P45bXO (Figure 6a).

**Figure 6.**
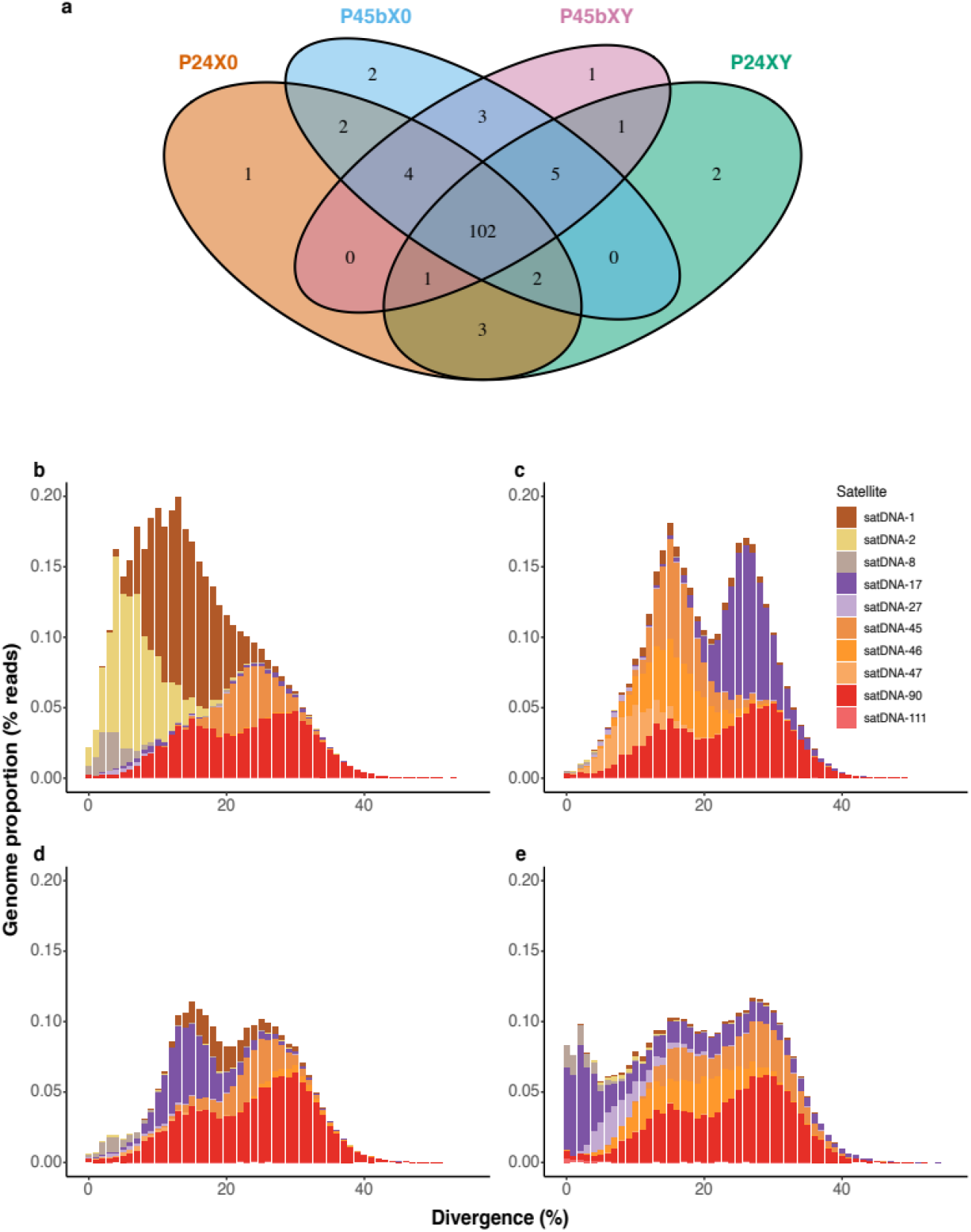
satDNA presence and divergence across four chromosomal races of the *viatica* species group. **a)** Venn diagram showing the number of satDNA families shared among the chromosomal races. **b)** Repeat landscapes based on RepeatMasker illustrating the abundance and divergence of 10 randomly selected satDNAs. **a)** P24X0, **b)** P24XY, **c)** P45bX0 and **d)** P45bXY. Color-coded bar plots showing the genome proportion (*y* axis) for each type of satDNAs in the sequenced reads analyzed, binned according to K2P distances to their corresponding consensus sequence (*x* axis, K2P value from 0 to 50 %). Copies in low-divergence bins are very similar to the consensus sequence and likely correspond to very recent copies. Out of the 102 satDNA families shared between all four races, there were 50 satDNA families showing differential amplification in copy number (cv > 80 %; Additional file 2: Table S8). From these 50 families, 10 randomly selected satDNAs are shown here for visualization of differential proliferation of satDNAs.

From the 102 satDNA families that were shared between all four races, there were 50 satDNA families that showed cv > 80 % (cv min 82 % and cv max 199 %) for abundance data between races (Additional file 1: Table S8), indicating differential amplification since the divergence of the chromosomal races. From these 50 satDNA families, we randomly selected 10 families to generate a repeat landscape of their relative genomic read abundance in 1 % bins of K2P distance. Differential amplification indicated by the differences in abundance and K2P distance distribution of sequences was significant for eight satDNAs (Kruskal-Wallis test *P* < 0.03; satDNA-1, satDNA-2, satDNA-8, satDNA-17, satDNA-27, satDNA-45, satDNA-47, satDNA-90) but not for the remaining two (*P* > 0.33; satDNA-46 and satDNA-111) (Figure 6, Table 5). Among these 10 satDNAs, the most abundant family was satDNA-1 in P24X0 (average 1.48 %) and the least abundant one was satDNA-47 in P24X0 (average K2P << 0.01 %). The most divergent family was satDNA-45 in P45bXY (average K2P 25.53 %) and the least divergent one was satDNA-8 in P45bXY (average K2P 4.01 %) (Table 5).

**Table 5:** Average of abundance and divergence of the 10 satDNAs shown in the Figure 6.

| Repeat | Abundance (%) |  |  |  | Divergence (% K2P) |  |  |  | Differential copy amplification<br>(Kruskal-Wallis test) |
| --- | --- | --- | --- | --- | --- | --- | --- | --- | --- |
|  | P24X0 | P24XY | P45bX0 | P45bXY | P24X0 | P24XY | P45bX0 | P45bXY |  |
| satDNA-1 | 1.48 | 0.13 | 0.23 | 0.06 | 13.94 | 19.60 | 20.28 | 19.86 | $P < 0.03$ |
| satDNA-2 | 0.96 | 0.01 | 0.01 | 0.02 | 7.12 | 9.50 | 11.31 | 10.18 | $P < 0.01$ |
| satDNA-8 | 0.15 | 0.01 | 0.06 | 0.08 | 5.10 | 4.72 | 6.57 | 4.01 | $P < 0.01$ |
| satDNA-17 | 0.06 | 0.50 | 0.57 | 0.75 | 24.4 | 15.77 | 19.48 | 19.90 | $P < 0.01$ |
| satDNA-27 | 0.02 | 0.03 | 0.03 | 0.19 | 6.73 | 9.51 | 8.44 | 9.29 | $P < 0.01$ |
| satDNA-45 | 0.39 | 0.53 | 0.32 | 0.51 | 25.33 | 17.38 | 23.58 | 25.53 | $P < 0.01$ |
| satDNA-46 | 0.01 | 0.51 | 0.05 | 0.44 | 16.64 | 16.42 | 17.12 | 18.30 | $P > 0.33$ |
| satDNA-47 | 0.01 | 0.27 | 0.03 | 0.01 | 11.57 | 10.01 | 9.83 | 11.14 | $P < 0.01$ |
| satDNA-90 | 1.20 | 1.05 | 1.20 | 1.20 | 22.82 | 23.02 | 23.85 | 23.67 | $P < 0.01$ |
| satDNA-111 | 0.004 | 0.01 | 0.005 | 0.02 | 22.46 | 15.34 | 22.84 | 9.78 | $P > 0.81$ |

## Discussion

### Genome assembly

The genomes of four chromosomal races of the *viatica* species group that we assembled here with 2.94-3.27 Gb assembly sizes are the third largest assembled insect genomes so far, with the largest being the two locust grasshoppers *L. migratoria* and *Schistocerca gregaria* with 6.5 and 8.6 Gb assembly sizes, respectively [50,51]. We believe that genome size estimates based on k-mer analysis better represent the genome size of these grasshoppers than the genome assembly sizes because highly repetitive sequences (e.g., centromeres, telomeres, satDNAs, non-recombining part oof sex chromosomes) are likely collapsed during the assembly process [61,62,73,74]. However, the two different k-mer approaches yielded quite different estimates between the chromosomal races (3.30-3.82 Gb by Supernova, and 5.42-6.32 Gb by findGSE), the reasons for which remain unclear. iThe large differences in contig size (contig N50 29.11-35.69 kb) of the assembled genomes of the *viatica* species group, *L. migratoria* (contig N50 9.30 kb) [50] and *S. gregaria* (contig N50 12.03 kb) [51] are probably due to the difference in the DNA library preparation and sequencing methods. The *L. migratoria* genome is based on Illumina short reads (2x 45-150 bp paired-end) while we used linked reads (150 bp paired-end Illumina short reads with barcode information from long input DNA molecules), and the *S. gregaria* assembly used paired-end and mate-pair Illumina short reads and PacBio long reads. The scaffold N50 (158 kb) of the *S. gregaria* genome assembly (8.6 Gb) is smaller than the scaffold N50 (317 kb) that we obtained in the P24XO race. In addition, the *S. gregaria* genome assembly is even more fragmented than the *L. migratoria* as indicated by the BUSCO scores (see [51]), indicating that assembling such large grasshopper genomes is challenging even using the combination of technologies above. Although the smaller genome assemblies of the *viatica* species group were more contiguous than the assembly of *L. migratoria*, they still contained missing and fragmented genes. This may be due to 1) large numbers of repetitive non-coding DNAs, 2) intron gigantism, 3) errors during the assembly process, 4) true missing genes, 5) failure in identifying any significant matches, and/or 6) failure in the gene prediction step to produce even a partial gene model that might have been recognized as a fragmented BUSCO match [60].

### TE dynamics and genome evolution

Approximately 66 to 72 % of the genome assembly of the *viatica* species group corresponds to TEs. These are even larger than the 60 to 62 % reported for the genome assemblies of *L. migratoria* [50] and *S. gregaria* [51] respectively, likely owing to the combination of methodological approaches to annotate TEs that we used. Given the deep divergence of Eumastacoidea and Acridoidea (~197 mya [33]), the two Orthopteran superfamilies to which the *viatica* species group and the locusts grasshoppers belong, respectively, these findings suggest that large repeatomes are widespread in grasshoppers. The *S. gregaria* genome assembly has 18,815 annotated genes, and the *L. migratoria* genome assembly has a similar number of annotated genes (17,307) to those recently reported in two cricket species with genome assembly sizes of 1.6 Gb (TE content 40 %; [75]), indicating the absence of partial or whole genome duplication events in these orthopteran lineages. Additionally, there was no evidence for such large-scale genome duplication events in the *viatica* species group because only 3-5 % of BUSCO genes were duplicated. Therefore, the large genome sizes in these grasshoppers are likely due to the expansion of TEs, which has been correlated with genome size evolution across the Tree of Life [2,76]. Indeed, we found massive recent amplification in hundreds of Mb (between 314 and 463 Mb) of the TE groups per genome assembly, an amount that is notably larger than the estimated genome size of many other insects [49,76]. The recent amplification mainly occurred in eight TE superfamilies (DNA/DNA, DNA/P, DNA/Sola, DNA/hAT, DNA/TcMar, RC/Helitron, LINE/L2, LTR/LTR, LTR/ Gypsy and SINE/tRNA) with largest variation for LTR/Gypsy and DNA/TcMar, indicating that the recent amplifications of TE superfamilies have widely shaped the TE landscape in the genomes of the chromosomal races. We thus suggest that the massive proliferation of TEs combined with a slow deletion rate might contribute to the genomic gigantism in grasshoppers, as proposed for other large eukaryotic genomes [77].

### Recently active TEs are permissively transcribed in gonads

We restricted our analysis to transcripts that originated from recent TEs (i.e., K2P < 5 %) because these are likely to be relevant source of transcriptional and transpositional activity. Our results demonstrated that recently active TE superfamilies from all five major TE groups (LINE/SINE/DNA/RC/LTR) are transcribed, suggesting that at least some of the transcribed TEs are capable of (retro)transposition. Recent TE expression tended to be differentially expressed in gonads compared to somatic tissues. This might indicate that TEs are transcribed and transpose themselves more frequently in gonads, transmitting new TE insertions to the next generation. The grasshopper ovaries and testes showed uneven expression of recent TEs, suggesting that substantial TE transcriptional variation likely exists across sexes. The expression variation of recent TEs between reproductive and somatic tissues is puzzling. We speculate that this might result from either global epigenetic reprograming during gametogenesis or the many more cell types/stages present in gonads than in individual somatic tissues of these grasshoppers. Alternatively, TE control might be tighter in somatic tissues, such that tight repression of TEs is important for the host and more feasible in somatic tissues (no global epigenetic reprogramming). It remains to be investigated if higher transcriptional activities of all five major TE groups, particularly in reproductive tissues, is associated with higher TE repressive mechanism activation (piwi/piRNA pathway [78–80]) to prevent the potentially deleterious effects of TE (retro)transposition in the host during global epigenetic reprogramming [78–81].

### The chromosomal races of *V. viatica* species group share a common collection of satDNAs which mostly experienced quantitative changes during evolution

To our knowledge, we uncovered the largest collection of satDNA families (129) ever reported for eukaryotic genomes. We propose that the 102 satDNA families shared among all four chromosomal races (Additional file 1: Table S6 and S8) represent the “library” present in the *viatica* ancestor. The 27 satDNA families that were not shared between all the four races either emerged after the divergence of the *viatica* ancestor or were lost in one or more races. This implies that the essential step in the evolution of a satDNA family might either be the acquisition of biological functions or the accumulation of sufficiently many copies to be maintained in the “library” over long evolutionary periods. How the novel satDNA families emerged remains unclear in the *viatica* species group, although unequal crossing-over, intra-strand homologous recombination, gene conversion, rolling-circle replication, and transposition are possible mechanisms [11,14–16,18,20,22]. After satDNA emergence, it is logical to assume that satDNA families stochastically expand or disappear and are only maintained in the long term if they acquire a function, such as in centromeres or heterochromatin formation. To test for satDNA functionality and to determine whether satDNAs were independently acquired or lost, additional data (e.g. chromosome *in situ* hybridization and ChIP-seq) is needed including races/species from the other morabine grasshoppers and under a robust phylogenetic hypothesis.

In line with the satDNA library hypothesis [23,24], 50 of the 102 satDNA families shared among all four chromosomal races experienced quantitative changes in copy number, and these happened over different K2P bins of divergence (see Figure 6b-d), suggesting rapid satDNA proliferation. The changes in copy number likely occurred by unequal crossing over, which is the mechanism that can yield changes in TR abundance, either as gains (amplifications) or losses (contractions) [82]. The large number of satDNA families in these chromosomal races is puzzling. The non-coding satDNAs have been traditionally viewed as mostly useless material capable of accumulating primarily in heterochromatin [11–13,27,29] until they become a too heavy load for the host genome (reviewed in [45]). It will be interesting to test whether the differential amplification of satDNAs is correlated with the amount of heterochromatin. On the other hand, it has been suggested that differential expression of satDNAs as satRNAs can cause genomic incompatibilities in hybrids because satRNAs play critical roles in kinetochore assembly (i.e. by binding to specific centromeric proteins like CENP-A and CENP-C) [25,26,45,46], heterochromatin formation [44,47,48] and function during cell division via siRNAs and piRNA pathways in *Schizosaccharomyces pombe, Drosophila*, nematodes, humans [44–48,83]. The satDNA families identified here are thus a set of candidates for future studies on which satDNAs are located in centromeres and which are involved in heterochromatin formation of grasshoppers.

### Conclusion

We have generated the so far most contiguous genome assemblies of grasshoppers with 66-75 % repeat contents using 10X Genomics linked-read sequencing. In-depth repeat annotation proved essential to elucidate the composition and characteristics of TEs and satDNAs in large and repetitive genomes such as grasshoppers. We showed a massive recent proliferation of a wide range of TEs, many of which are transcribed more frequently in germline than somatic tissues. In addition, we uncovered the largest number of satDNA families ever reported in eukaryotic genomes, and showed that, despite the recent divergence of the four *viatica* chromosomal races, satDNA evolution underwent rapid expansions or contractions in copy number.

## Methods

### Taxon sampling, DNA and RNA extraction, and sequencing

Chromosomal races of *Vandiemenella* morabine grasshoppers were collected between 2002 and 2017 in South Australia. To identify the races, the testes were dissected from males and fixed for karyotyping as described previously [53,84]. The remaining body parts were flash-frozen in liquid nitrogen and stored at −80 °C in the Australian Biological Tissue Collection until subsequent DNA extraction. One male per race (P24X0, P24XY, P45bX0 and P45bXY) was used for DNA extraction from either heads or legs using the MagAttract HMW DNA Kit (QIAGEN, Hilden, Germany; Cat No. 67563). Sequencing libraries were prepared as recommended by Chromium Genome preparation kit (10X Genomics, Inc., Pleasanton, CA, USA; Cat No. 120215). Paired-end read sequencing (2×150 bp) was performed on the Illumina HiSeq X (Illumina, Inc., San Diego, CA, USA) using the Chromium library.

Tissues from males (head, leg, and testes) and females (head, leg, and ovary) of the chromosomal races P24X0 and P24XY were dissected and placed in RNAlater (ThermoFisher Scientific, Waltham, MA, USA) and stored at −80 °C until subsequent RNA extraction. In total, 42 males and 62 females were used for RNA extraction (5-11 individuals per tissue/sex/race). We extracted RNA with phenol-based phase separation using the TRIzol reagent (ThermoFisher Scientific) following the standard protocol recommended by the supplier. Sequencing libraries were prepared according to the TruSeq stranded mRNA library preparation kit (Illumina, Inc., Cat No.20020594/5) including poly-A selection. Paired-reads (2×100 bp) were sequenced on the NovaSeq 6000 S2 flowcell (Illumina, Inc.). DNA and RNA library preparation and sequencing were performed at the SNP&SEQ Technology Platform in Uppsala, Uppsala University, Department of Medical Sciences, Uppsala Biomedical Centre (BMC) (Uppsala, Sweden).

### Genome size estimation and genome assembly

We performed genome size estimation by counting k-mer frequency of the quality checked 10X Genomics linked-reads. The multiple fastq.gz files were two-step processed for counting k-mers using Jellyfish 2.2.6 [85] with the following setting: -t 8 -C -m 18 -s 5G --min-quality=20 -- quality-start=33. We used the R package findGSE [63] to estimate genome sizes by using the output of Jellyfish.

We used Supernova 2.1.0 [59] to generate a “pseudohaploid” genome assembly of males of P24X0, P24XY, P45bX0 and P45bXY using 10X Genomics linked-read data. Supernova uses the H0X Genomics linked reads generated from a single library of DNA from an individual organism as source, potentially allowing for the assembly of longer contigs and scaffolds than conventional short-read technologies [59]. For comparative analysis of the generated draft assemblies, we downloaded the *Locusta migratoria* genome from the NCBI database (GCA_000516895.1) [50]. The completeness of the assemblies was evaluated with BUSCO v3 [60] with the Arthropoda database [86] as a reference. The recovered matches were classified as complete if their lengths were within the expectation of the BUSCO profile match lengths. If these were found more than once, they were classified as duplicated. The matches that were only partially recovered were classified as fragmented, and BUSCO groups that passed the test of gene prediction but for which there were no matches in the database were classified as missing.

### Assessing the TEs content across genomes

We used the draft male genome assemblies of P24X0, P24XY, P45bX0 and P45bXY to generate repeat libraries for each of the four genomes using RepeatModeler 1.0.8 [64]. Because the P24X0 had the highest assembly contiguity and BUSCO scores, the repeats classified as unknown in the P24X0 RepeatModeler library were selected for manual curation following the method used in Suh et al. [87]. Every consensus sequence was aligned back to the assembled genome sequence of P24X0, then the best 20 BLASTn hits were collected, extended by 2 kb and aligned to one another using MAFFT 7 [88]. Manually curated consensus sequences of P24X0 were then classified as TE families/subfamilies based on the proposed classification system for TEs (open reading frames, terminal repeats, target site duplications) [7] and Repbase similarity searches [89]. We removed redundancies from the curated TE consensus sequences of P24X0 by merging sequences that were greater than 80% similar using CD-HIT-EST [90], with the following setting: -c 0.80 -n 5-M 0 -aS 0.80 -G 0 -g 1.

The curated TE library from P24X0 race was used as a reference to re-classify repeats in the TE libraries of the other three races (i.e., P24XY, P45bX0 and P45bXY) by using RepeatMasker 4.0.8 [65]. We removed redundancies of the re-classified TEs within each library using CD-HIT-EST. We thus obtained the following combinations of libraries: 1) P24X0 RepeatModeler library (RML) + Arthropoda Repbase library (ARL) + the curated library, 2) P24XY RML + ARL + curated library, 3) P45bX0 RML + ARL + curated library, and 4) P45XY RML + ARL + curated library. We used these libraries to mask their respective genomes to annotate repeats using RepeatMasker [65]. We then used each of the .align RepeatMasker output files per genome to estimate abundance and divergence of each TE superfamily. To visualize the temporal activity/accumulation of TEs across races, we generated landscape bar plots depicting the relative abundance of repeat elements on the *y* axis and the K2P distance from the respective consensus sequence on the *x* axis.

In addition to the annotation of TEs in the four repeat libraries above, we also searched for new Helitrons across the genome assemblies using HelitronScanner 1.0 [66]. HelitronScanner uses the two-layered local combinational variable (LCV) tool for Helitron identification. To avoid false positive in Helitron searches, we used thresholds of 7 for both Helitron ends (14 for sum of LCV scores) as a parameter in the HelitronScanner runs. We used CD-HIT-EST to remove redundancies of the detected Helitron sequences by merging sequences that were greater than 80 % similar. This produced a set of non-redundant representative sequences (families) of new Helitrons. We used the detected Helitrons in each chromosomal race in combination with the libraries above mentioned to finally mask the respective genomes.

We further quantified the repeatome between races using read-based approaches to compare with the assembly-based approach. For this purpose, the relative genomic abundance of repeats in the sequenced reads were compared by random sampling of 4 million read pairs per library and masking them with the combined repeat libraries (RepeatModeler + HelitronScanner + TRs) using RepeatMasker. This quantification was done separately in each race using their respective repeat library. TRs were detected using RepeatExplorer2 [67–69] and NOVOplasty 3.7.2 [70] (see below).

### Transcription of TEs

We used Illumina RNA-seq reads (2x 100 bp) from males and females (testis, ovary, head and leg) of P24X0 and P24XY races to investigate the transcriptional profile of TEs in each tissue. For comparative purposes, we used the RepeatMasker annotation (with simple and low complexity repeats removed) from the P24X0 and P24XY races to check whether the TEs show differential expression among sexes and tissues, and to compare the intensity and direction of the bias among races. RNA-seq reads per sample were trimmed with TrimGalore (https://github.com/FelixKrueger/TrimGalore). Erroneous k-mers from Illumina paired-end reads were removed using rCorrector [91]. Unfixable reads (often riddled with N nucleotides or represented by other low complexity sequences) were discarded using FilterUncorrectablePEfsta.py script obtained from the Harvard Informatics GitHub repository (https://github.com/harvardinformatics/TranscriptomeAssemblyTools). We used SortMeRNA 3.0.3 [92] for local alignment, filtering, mapping, and clustering to remove rRNA.

We applied the RepEnrich2 protocol (https://github.com/nerettilab/RepEnrich2) to estimate TE transcript expression levels in each genome assembly. To assign mapped reads to a genomic locus, RepEnrich2 requires an annotation file that specifies repeat element coordinates. The RepeatMasker annotation was used for the approach. We estimated the expression levels of reads that originated from recent TE copies. For this analysis, we provided RepEnrich2 with a filtered RepeatMasker annotation containing only TE copies with < 5 % K2P from the consensus sequences. The filtered RepeatMasker annotation was then used to build the final annotation required for RepEnrich2 protocol by running the RepEnrich2_setup script. Next, the pre-processed RNA-seq reads were mapped to each of the assemblies using Bowtie2 2.2.9 [93] as the read mapper. The quantification step included raw read count and differential expression analysis of TEs estimated by the RepEnrich2 protocol and DESeq2 1.20.0 [72], respectively. The data regarding sex-biased TE expression were compared between gonads, heads, and legs from males and females of the chromosomal races P24X0 and P24XY. The statistical analysis of differentially expressed TE was performed using DESeq2 as implemented in the Bioconductor package [94] in R [95]. All *P*-values were adjusted using the Wald test as implemented in DESeq2. A TE was considered biased if the comparison for the factor condition (samples) yielded an adjusted *P* < 0.05. The degree of bias was determined by the log2 fold-change (log2FC) difference between conditions as calculated in DESeq2, i.e., those TEs with log2FC > 0 and log2FC < 0 and with an adjusted *P* < 0.05 were considered as male-biased TE (MBTEs) and female-biased TE (FBTEs), respectively.

### Tandem repeat detection in sequencing reads

We used RepeatExplorer2 [67–69] to identify satDNAs using the linked reads of each chromosomal race used for genome assembly above. RepeatExplorer was run separately in each race. Prior to RepeatExplorer2 graph-based clustering analysis, sequence reads were quality-trimmed using TrimGalore. The trimmed paired-end reads were joined by using the “fastq-join” software of the FASTX-Toolkit suit [96] with default parameters. The joined paired-end reads were then subject to graph-based clustering and assembly using RepeatExplorer2. A set of randomly selected 2,000,000 Chromium linked reads with average length of 150 bp were used as an input for clustering analysis.

We used the dotplot graphic alignment tool implemented in Dotlet [97] to confirm the tandem organization of those clusters with high graph density in RepeatExplorer2 output. Additionally, we used the FlexiDot 1.06 [98] suite to generate all-against-all dotplots visualization of those clusters identified as tandem arrays. The monomer with maximum length was used as the representative copy for each satDNA family, and as the query sequences in further BLASTn (http://www.ncbi.nlm.gov/Blast/) and Repbase [89] searches to check for similarity with published sequences, and as a query to check for overlap with the RepeatModeler libraries. We named each satDNA family as Vv (from *Vandiemenella viatica* group) following by the provisional taxon name (i.e., P24X0, P24XY, P45bX0 and P45bXY) and a number in descending order of the genome proportion. To search for HORs, we counted the maximum number of tandem monomer arrays per contig for each satDNA family by analyzing the dotplots. We then counted the total number of monomers present in each cluster. The multiple sequence alignments of satDNA copies were generated using Muscle [96] implemented in MEGA 5 [99].

We ran NOVOPlasty 3.7.2 [70] to search for tandem multigene families, such as 45S rRNA, 5S rRNA, U snRNA (U1, U2, U5, U6), and histone genes (H1, H2A, H2B, H3 and H4). We used known seed sequences of these genes coming from other species and k-mer between 21 and 23 as parameters for assembly: Gomphocerinae sp. for the 45S rRNA gene (AY859546.1), *Ronderosia bergi* for the 5S rRNA gene (KP213274), *Nasonia vitripennis* for the H1 histone gene (XM_003423983.3), *Culex quinquefasciatus* for the H2A (XM_001862631.1) and H2B (XM_001870471.1) histone genes, *Locusta migratoria* for the H3 histone gene (GU111931.1), *N. vitripennis* for the H4 histone gene (XM_001599180.3:49-357), *Eyprepocnemis plorans* for the U1 snRNA gene (KJ606069.1), *Abracris flavolineata* for the U2 rRNA gene (KP975085.1), and *D. melanogaster* for the U5 snRNA (NR_001933.1) and U6 snRNA genes (NR_002081.1). The canonical tandem telomere repeat (TTAGG)_n_ recognized in insects [100] was recovered from the RepeatExplorer2 output.

The relative genomic abundances and K2P divergences of TRs were estimated individually in each race by sampling 4 million read pairs per library retrieved by seqtk (https://github.com/lh3/seqtk) and aligning them to the TR database with RepeatMasker. The sampled reads were mapped to dimers of satDNA consensus sequences, and for smaller TRs, several monomers were concatenated until reaching roughly 150 bp array length. To visualize the temporal accumulation of TRs across races, we generated TR landscape plots that depict the relative abundance of repeat elements on the *y* axis and the K2P distance from the respective consensus sequence on the *x* axis.

To search for homologous satDNAs between races, we used two approaches. First, we built a satDNA database by concatenating all the satDNA consensus sequences detected in each race and performed an all-against-all comparison of the consensus sequences in the database using the rm.homology.py script (https://github.com/fjruizruano/ngs-protocols) [31]. Second, we mapped the genomic reads of each race separately to the satDNA consensus database using RepeatMasker, as (mentioned above, and recorded presence/absence of satDNAs in each of the four races.

## Abbreviations

A+T%: Percentage of A + T bases, i.e., the molar ratio of adenine and thymine in DNA
cv: Coefficient of variation
HOR: Higher-order repeat structure
K2P: Kimura 2-parameter distance
LINE: Long interspersed element
LTR: Long terminal repeat
SINE: Short interspersed element
TE: Transposable element
TIR: Terminal inverted repeat
TR: Tandem repeat
RC: Rolling-circle transposable element
satDNA: satellite DNA
MBTE: Male-biased transposable element
FBTE: Female-biased transposable element
log2FC: log2 fold-change

## Availability of data and materials

The datasets generated as part of the study are available as supplementary information. Repeat consensus sequences are available as Additional file 3 (P24X0), Additional file 4 (P24XY), Additional file 5 (P45bX0) and Additional file 6 (P45bXY). Each of these additional files contain (libraries generated by RepeatModeler (including curated and re-classified repeats), HelitronScanner and RepeatExplorer2. Raw reads and assemblies were deposited in Sequence Read Archive (accession numbers XXXXXXXX-XXXXXXXX)..

## Author’s contribution

Conceptualization: OMPG, TK, AS; Formal analysis: OMPG, JK; Funding acquisition: OMPG, TK, AS; Investigation: OMPG, JK, MPF; Project Administration: TK, AS; Resources: TK, AS, TMB, SJBC, KKJ; Supervision: TK, AS; Validation: OMPG, JK; Visualization: OMPG; Writing – original draft: OMPG; Writing – remaining drafts: OMPG, TK, AS, TMB, SJBC, KKJ.

## Acknowledgments

The manuscript was improved by comments from the Julie Blommaert, Valentina Peona, Francisco J. Ruiz-Ruano and Diogo C. Cabral-de-Mello. TK is grateful for Roger Butlin and the University of Sheffield for supporting this research during the INCA fellowship. Computations (were performed on resources provided by the Swedish National Infrastructure for Computing (SNIC) through the Uppsala Multidisciplinary Center for Advanced Computational Science (UPPMAX). Sequencing was performed by the SNP&SEQ Technology Platform in Uppsala. The facility is part of the National Genomics Infrastructure (NGI) Sweden and Science for Life Laboratory. The SNP&SEQ Platform is also supported by the Swedish Research Council and the Knut and Alice Wallenberg Foundation.

