## Additional file 1 for "Too much too many: comparative analysis of morabine grasshopper genomes reveals highly abundant transposable elements and rapidly proliferating satellite DNA repeats"

**Table S1:** Total numbers of differentially expressed TE subfamilies (consensus sequences) containing all TEs between sexes and tissues in two chromosomal races of the *viatica* species group.

| Race | Sample | No. of expressed TEs | SBTEs (%) | MBTEs (%) | FBTEs (%) |
|---|---|---|---|---|---|
| P24XO | Head | 2,354 | 18 (0.76) | 12 (0.51) | 6 (0.25) |
| | Gonad | 2,778 | 809 (29.12) | 441 (16) | 368 (13) |
| | Leg | 2,629 | 1 (0.04) | 1 (0.04) | 0 |
| P24XY | Head | 7,240 | 36 (0.49) | 29 (0.40) | 7 (0.09) |
| | Gonad | 7,240 | 454 (6.30) | 377 (5.20) | 77 (1.10) |
| | Leg | 7,240 | 171 (2.36) | 166 (2.30) | 31 (0.43) |

We report the total number of TEs expressed, and those TEs that were sex-biased with log2FC > 0 and log2FC < 0 and adjusted $P < 0.05$; log2FC = log2 fold-change. SBTEs = sex-biased TEs; MBTE = male-biased TEs.

**Table S2:** Most abundant tandem repeats (TRs) in the P24XO sex chromosome race (male reads). The repeats are sorted in descending order of the genomic read proportion.

| Repeat name | Consensus length (bp) | Genome proportion % | A+T % | K2P divergence % | Total of monomers in clusters | Max. number of monomers per tandem arrays per contig | Total repeat family length (kb) | HORs |
|---|---|---|---|---|---|---|---|---|
| VvP24XO-1 | 51 | 1.48 | 52.9 | 13.94 | 182 | 9 | 9.282 | Yes |
| VvP24XO-2 | 53 | 0.96 | 64.2 | 7.12 | 50 | 8 | 2.65 | Yes |
| VvP24XO-3 | 257 | 0.94 | 64.6 | 5.24 | 10 | 3 | 2.57 | No |
| VvP24XO-4 | 53 | 0.89 | 58.5 | 6.3 | 50 | 8 | 2.65 | Yes |
| VvP24XO-5 | 189 | 0.74 | 52.9 | 21.1 | 61 | 3 | 11.529 | Yes |
| Telomere repeat | 5 | 0.45 | 60 | 1.92 | - | - | - | No |
| VvP24XO-6 | 7 | 0.32 | 71.4 | 14.02 | 2327 | 60 | 65.156 | Yes |
| VvP24XO-7 | 147 | 0.31 | 64.6 | 19.96 | 103 | 3 | 13.377 | Yes |
| VvP24XO-7 | 66 | 0.17 | 59.1 | 8.11 | 48 | 6 | 3.168 | Yes |
| VvP24XO-9 | 34 | 0.15 | 61.8 | 5.1 | 50 | 10 | 1.7 | Yes |
| 45S rRNA | 3432 | 0.13 | 42.2 | 1.62 | - | - | 3432 | No |
| VvP24XO-10 | 21 | 0.09 | 52.4 | 7.8 | 541 | 22 | 11.361 | Yes |
| VvP24XO-11 | 125 | 0.07 | 60.8 | 5.31 | 45 | 3 | 5.625 | Yes |
| VvP24XO-12 | 80 | 0.07 | 53.7 | 6.5 | 25 | 5 | 2.015 | Yes |
| VvP24XO-13 | 1140 | 0.07 | 54 | 3.16 | 35 | 2 | 39.9 | Yes |
| VvP24XO-14 | 101 | 0.07 | 55.4 | 4.44 | 16 | 4 | 1.616 | No |
| VvP24XO-15 | 120 | 0.07 | 55 | 10.1 | 72 | 4 | 8.64 | Yes |
| VvP24XO-16 | 75 | 0.07 | 46.7 | 7.27 | 65 | 5 | 4.875 | Yes |
| VvP24XO-17 | 130 | 0.06 | 62.3 | 5.4 | 14 | 3 | 1.82 | No |
| VvP24XO-18 | 51 | 0.06 | 48.5 | 24.4 | 50 | 6 | 5.05 | Yes |

| | | | | | | | | |
|---|---|---|---|---|---|---|---|---|
| VvP24XO-19 | 31 | 0.06 | 54.8 | 6.91 | 90 | 12 | 2.79 | Yes |
| VvP24XO-20 | 133 | 0.05 | 74.4 | 8.22 | 39 | 3 | 5.187 | No |
| VvP24XO-21 | 52 | 0.04 | 57.7 | 6.46 | 43 | 8 | 2.236 | Yes |
| VvP24XO-22 | 54 | 0.04 | 55.6 | 5.42 | 35 | 7 | 1.89 | Yes |
| VvP24XO-23 | 66 | 0.04 | 51.5 | 7.9 | 302 | 9 | 19.932 | Yes |
| VvP24XO-24 | 32 | 0.03 | 65.7 | 6.67 | 214 | 12 | 6.848 | Yes |
| VvP24XO-25 | 52 | 0.03 | 50 | 8.01 | 87 | 9 | 4.524 | Yes |
| VvP24XO-26 | 220 | 0.03 | 63.2 | 10.71 | 5 | 2 | 1.1 | No |
| VvP24XO-27 | 79 | 0.03 | 59.5 | 7.43 | 62 | 6 | 4.898 | Yes |
| VvP24XO-28 | 52 | 0.02 | 46.2 | 6.73 | 127 | 9 | 6.604 | Yes |
| VvP24XO-29 | 26 | 0.02 | 56.4 | 11.73 | 60 | 13 | 4.68 | Yes |
| VvP24XO-30 | 131 | 0.02 | 61.1 | 7.82 | 39 | 3 | 5.109 | Yes |
| VvP24XO-31 | 63 | 0.02 | 69.8 | 7.94 | 30 | 6 | 1.89 | Yes |
| VvP24XO-32 | 122 | 0.02 | 38.5 | 13.66 | 47 | 3 | 5.734 | Yes |
| VvP24XO-33 | 171 | 0.02 | 63.7 | 8.36 | 66 | 3 | 11.286 | Yes |
| VvP24XO-34 | 122 | 0.02 | 62.3 | 5.92 | 58 | 4 | 7.076 | Yes |
| 5S rRNA + NTS | 300 | 0.02 | 51.7 | 8.5 | - | - | - | No |
| VvP24XO-35 | 42 | 0.01 | 50 | 7.56 | 300 | 18 | 12.6 | Yes |
| VvP24XO-36 | 60 | 0.01 | 45 | 16.72 | 98 | 8 | 5.88 | Yes |
| VvP24XO-37 | 108 | 0.01 | 59.3 | 3.99 | 20 | 4 | 2.16 | Yes |
| VvP24XO-38 | 82 | 0.01 | 59.8 | 3.36 | 25 | 5 | 2.05 | Yes |
| VvP24XO-39 | 138 | 0.01 | 65.2 | 6.7 | 15 | 3 | 2.78 | Yes |
| VvP24XO-40 | 86 | 0.01 | 52.3 | 4.02 | 20 | 4 | 1.72 | Yes |
| VvP24XO-41 | 26 | 0.01 | 38.5 | 6.91 | 197 | 16 | 5.122 | Yes |
| VvP24XO-42 | 33 | 0.01 | 46.3 | 10.17 | 224 | 12 | 7.392 | Yes |

| | | | | | | | | |
|---|---|---|---|---|---|---|---|---|
| VvP24XO-43 | 76 | 0.01 | 44.7 | 6.8 | 75 | 6 | 5.7 | Yes |
| VvP24XO-44 | 36 | 0.01 | 55.6 | 11.84 | 120 | 12 | 4.32 | Yes |
| VvP24XO-45 | 65 | 0.01 | 66.2 | 3.68 | 30 | 6 | 1.95 | Yes |
| H2A histone | 360 | 0.01 | 41.9 | 1.43 | - | - | - | - |
| H4 histone | 361 | 0.0047 | 44 | 2.31 | - | - | - | - |
| H2B histone | 601 | 0.0044 | 44.4 | 0.97 | - | - | - | - |
| H3 histone | 331 | 0.0044 | 39.1 | 0.55 | - | - | - | - |
| U2 snRNA | 178 | 0.0038 | 53.9 | 3.81 | - | - | - | - |
| U1 snRNA | 162 | 0.0015 | 45.1 | 2.37 | - | - | - | - |
| U5 snRNA | 123 | 0.001 | 57.7 | 4.45 | - | - | - | - |
| U6 snRNA | 111 | 0.00024 | 55.9 | 4.62 | - | - | - | - |

The satDNA families and the telomere tandem repeat were assembled by RepeatExplorer2, and the tandem multigene families were assembled by NOVOPlasty. Abundance and divergence were estimated by RepeatMasker. satDNA families are named as VvP24XO; with "Vv" for *Vandiemenella viatica* group and "P" for provisional taxon name, and a number in a descending order of the genomic read proportion. A+T %: percentage of A+T in the consensus sequences. K2P divergence %: Kimura 2-parameter divergence from the consensus sequence. Total of monomers in clusters: number of monomers recovered for each satDNA family in each the clusters in RepeatExplorer output. Max. number of monomers per tandem arrays per contigs: maximum number of tandemly repeated monomers found in the largest contig within the cluster. Total repeat family length (kb): length of each satDNA family in kb obtained by the total of monomers in clusters per consensus size. HORs: High order repeat structures identified by visual inspection in a dotplot.

**Table S3:** Most abundant tandem repeats (TRs) in the P24XY sex chromosome race (male reads). The repeats are sorted in descending order of the male genome proportion.

| **Repeat name** | **Consensus length (bp)** | **Genome proportion %** | **A+T %** | **K2P divergence %** | **Total of monomers in clusters** | **Max. number of monomers per tandem arrays per contig** | **Total repeat family length (kb)** | **HORs** |
|---|---|---|---|---|---|---|---|---|
| Telomere | 5 | 1.59 | 40 | 2.6 | 690 | 66 | 3.45 | - |
| VvP24XY-1 | 189 | 0.88 | 56 | 11.96 | 15 | 2 | 5.52 | Yes |
| VvP24XY-2 | 52 | 0.57 | 50 | 19.48 | 110 | 8 | 5.72 | Yes |
| VvP24XY-3 | 50 | 0.53 | 58 | 17.38 | 72 | 8 | 4.176 | Yes |
| VvP24XY-4 | 168 | 0.51 | 67.9 | 16.42 | 177 | 3 | 29.736 | Yes |
| VvP24XY-5 | 19 | 0.27 | 63.2 | 10.01 | 1054 | 24 | 20.026 | Yes |
| VvP24XY-6 | 147 | 0.26 | 64.6 | 18.77 | 245 | 3 | 51.156 | Yes |
| VvP24XY-7 | 7 | 0.19 | 71.4 | 13.65 | 1741 | 60 | 12.187 | Yes |
| VvP24XY-8 | 51 | 0.13 | 58.5 | 19.6 | 341 | 10 | 17.391 | Yes |
| 45S rRNA | 5693 | 0.11 | 42.8 | 0.88 | - | - | - | - |
| VvP24XY-9 | 80 | 0.08 | 52.5 | 7.23 | 37 | 6 | 2.96 | Yes |
| VvP24XY-10 | 130 | 0.05 | 60.8 | 6.58 | 21 | 3 | 2.73 | Yes |
| VvP24XY-11 | 133 | 0.05 | 74.4 | 5.23 | 33 | 3 | 5.187 | Yes |
| VvP24XY-12 | 43 | 0.05 | 48.8 | 16.18 | 206 | 9 | 8.858 | Yes |
| VvP24XY-13 | 75 | 0.05 | 46.7 | 7.35 | 65 | 5 | 4.875 | Yes |
| VvP24XY-14 | 130 | 0.05 | 63.1 | 5.41 | 27 | 3 | 3.51 | Yes |
| VvP24XY-15 | 52 | 0.04 | 55.8 | 7.36 | 43 | 8 | 2.236 | Yes |
| VvP24XY-16 | 31 | 0.04 | 54.8 | 6.85 | 91 | 12 | 2.821 | Yes |
| VvP24XY-17 | 52 | 0.04 | 46.2 | 5.66 | 49 | 7 | 2.548 | Yes |
| VvP24XY-18 | 220 | 0.04 | 62.7 | 7.72 | 7 | 2 | 1.54 | Yes |
| VvP24XY-19 | 30 | 0.03 | 50 | 5.58 | 136 | 12 | 4.08 | Yes |

| | | | | | | | | |
|---|---|---|---|---|---|---|---|---|
| VvP24XY-20 | 53 | 0.03 | 45.3 | 9.51 | 128 | 9 | 6.784 | Yes |
| VvP24XY-21 | 43 | 0.03 | 51.2 | 17.17 | 135 | 8 | 5.805 | Yes |
| VvP24XY-22 | 61 | 0.03 | 67.2 | 8.47 | 48 | 6 | 2.928 | Yes |
| VvP24XY-23 | 26 | 0.03 | 53.8 | 9.43 | 167 | 15 | 4.342 | Yes |
| VvP24XY-24 | 32 | 0.02 | 65.6 | 7.1 | 272 | 12 | 8.704 | Yes |
| VvP24XY-25 | 1070 | 0.02 | 56 | 7.92 | 20 | 1 | 21.4 | Yes |
| VvP24XY-26 | 68 | 0.02 | 55.9 | 5.46 | 25 | 5 | 1.7 | Yes |
| VvP24XY-27 | 79 | 0.02 | 59.5 | 7.31 | 42 | 7 | 3.318 | Yes |
| VvP24XY-28 | 66 | 0.02 | 56.1 | 11.24 | 134 | 9 | 8.944 | Yes |
| VvP24XY-29 | 131 | 0.02 | 61.1 | 7.47 | 33 | 3 | 4.323 | Yes |
| VvP24XY-30 | 171 | 0.02 | 63.5 | 6.45 | 24 | 3 | 4.104 | Yes |
| VvP24XY-31 | 1690 | 0.02 | 59.3 | 4.84 | 15 | 1 | 25.35 | Yes |
| VvP24XY-32 | 123 | 0.02 | 61 | 5.74 | 39 | 3 | 4.797 | Yes |
| VvP24XY-33 | 138 | 0.02 | 65.2 | 3.69 | 27 | 3 | 3.726 | Yes |
| VvP24XY-34 | 63 | 0.01 | 69.8 | 4.27 | 48 | 6 | 3.024 | Yes |
| VvP24XY-35 | 20 | 0.01 | 65 | 10.71 | 765 | 27 | 15.3 | Yes |
| VvP24XY-36 | 121 | 0.01 | 59.5 | 6.36 | 85 | 4 | 10.285 | Yes |
| VvP24XY-37 | 60 | 0.01 | 43.3 | 12.29 | 44 | 8 | 2.64 | Yes |
| VvP24XY-38 | 54 | 0.01 | 55.6 | 5.85 | 43 | 8 | 2.322 | Yes |
| U2 snRNA | 213 | 0.01 | 54.5 | 1.67 | - | - | - | - |
| VvP24XY-39 | 36 | 0.01 | 47.2 | 9.92 | 144 | 12 | 5.184 | Yes |
| VvP24XY-40 | 104 | 0.01 | 53.8 | 9.46 | 81 | 4 | 8.424 | Yes |
| VvP24XY-41 | 29 | 0.01 | 41.4 | 6.18 | 131 | 13 | 3.799 | Yes |
| VvP24XY-42 | 34 | 0.01 | 58.8 | 4.72 | 60 | 10 | 2.04 | Yes |
| VvP24XY-43 | 32 | 0.01 | 59.4 | 22.67 | 392 | 16 | 12.544 | Yes |
| VvP24XY-44 | 108 | 0.01 | 59.3 | 3.15 | 20 | 4 | 2.16 | Yes |
| VvP24XY-45 | 42 | 0.01 | 48.8 | 6.86 | 297 | 12 | 12.474 | Yes |

| | | | | | | | | |
|---|---|---|---|---|---|---|---|---|
| VvP24XY-46 | 15 | 0.01 | 53.3 | 2.88 | 250 | 22 | 3.75 | Yes |
| VvP24XY-47 | 19 | 0.01 | 47.4 | 7.33 | 139 | 18 | 2.641 | Yes |
| VvP24XY-48 | 42 | 0.01 | 59.5 | 4.63 | 40 | 8 | 1.68 | Yes |
| H3 histone | 412 | 0.0047 | 40.5 | 1.46 | - | - | - | - |
| H2B histone | 428 | 0.0046 | 47.1 | 1.29 | - | - | - | - |
| H1 histone | 347 | 0.0046 | 39.9 | 1.86 | - | - | - | - |
| H2A histone | 383 | 0.0043 | 42.7 | 1.12 | - | - | - | - |
| H4 histone | 201 | 0.0029 | 46.8 | 1.04 | - | - | - | - |
| U1 snRNA | 162 | 0.0015 | 44.4 | 2.4 | - | - | - | - |
| U5 snRNA | 117 | 0.0011 | 58.1 | 1.46 | - | - | - | - |
| U6 snRNA | 107 | 0.00042 | 56.1 | 5.36 | - | - | - | - |
| 5S rRNA | 119 | 0.0004 | 37 | 8.7 | - | - | - | - |

The satDNA families and the telomere tandem repeat were assembled by RepeatExplorer2, and the tandem multigene families were assembled by NOVOPlasty. Abundance and divergence were estimated by RepeatMasker. satDNA families are named as VvP24XY; with "Vv" for *Vandiemenella viatica* group following by the "P" from provisional taxon name, and a number in a descending order of the male genomic read proportion. A+T %: percentage of A+T in the consensus sequences. Kimura divergence %: Kimura 2-parameter divergence from the consensus sequence. Total of monomers in clusters: number of monomers recovered for each satDNA family in each the clusters in RepeatExplorer output. Max. number of monomers per tandem arrays per contig: maximum number of tandemly repeated monomers found in the largest contig within the cluster. Total repeat family length (kb): length of each satDNA family in kb obtained by the total of monomers in cluster per consensus size. HORs: High order repeat structures identified by visual inspection in a dotplot.

**Table S4:** Most abundant tandem repeats (TRs) in the P45bXO sex chromosome (male reads). The repeats are sorted in descending order of the genome proportion.

| Repeat name | Consensus length (bp) | Genome proportion % | A+T % | K2P divergence % | Total of monomers in clusters | Max. number of monomers per tandem arrays per contig | Total repeat family length (kb) | HORs |
|---|---|---|---|---|---|---|---|---|
| Telomere repeat | 5 | 0.94 | 60 | 3.35 | - | - | - | - |
| VvP45bXO-1 | 1700 | 0.89 | 58.9 | 21.68 | 112 | 1 | 190.4 | Yes |
| VvP45bXO-2 | 189 | 0.77 | 56 | 12.79 | 32 | 2 | 11.776 | No |
| 45S rRNA | 5758 | 0.61 | 45.4 | 4.35 | - | - | - | - |
| VvP45bXO-3 | 51 | 0.50 | 52.9 | 15.77 | 163 | 9 | 8.313 | Yes |
| VvP45bXO-4 | 21 | 0.46 | 52.4 | 8.05 | 396 | 20 | 8.316 | Yes |
| VvP45bXO-5 | 282 | 0.44 | 64.8 | 19.87 | 185 | 3 | 52.17 | No |
| VvP45bXO-6 | 101 | 0.38 | 51.5 | 8.53 | 56 | 4 | 5.656 | Yes |
| VvP45bXO-7 | 51 | 0.23 | 51 | 20.28 | 168 | 8 | 8.568 | Yes |
| VvP45bXO-8 | 80 | 0.21 | 55 | 8.31 | 46 | 6 | 3.68 | Yes |
| VvP45bXO-9 | 147 | 0.18 | 64.6 | 13.79 | 157 | 3 | 15.435 | Yes |
| VvP45bXO-10 | 54 | 0.16 | 51.9 | 8.93 | 91 | 9 | 4.914 | Yes |
| VvP45bXO-11 | 20 | 0.13 | 65 | 12.89 | 236 | 10 | 9.44 | Yes |
| VvP45bXO-12 | 63 | 0.10 | 68.3 | 4.89 | 64 | 6 | 3.402 | Yes |
| VvP45bXO-13 | 99 | 0.09 | 61.6 | 10.25 | 60 | 6 | 5.94 | Yes |
| VvP45bXO-14 | 67 | 0.08 | 58.2 | 6.16 | 25 | 5 | 1.675 | Yes |
| VvP45bXO-15 | 52 | 0.07 | 63.8 | 7.07 | 43 | 8 | 2.236 | Yes |
| VvP45bXO-16 | 130 | 0.06 | 59.2 | 5.96 | 27 | 3 | 3.51 | Yes |
| VvP45bXO-17 | 34 | 0.06 | 61.8 | 6.57 | 82 | 12 | 2.788 | Yes |
| VvP45bXO-18 | 32 | 0.06 | 61.6 | 6.73 | 226 | 12 | 7.242 | Yes |
| VvP45bXO-19 | 16 | 0.06 | 50 | 8.49 | 414 | 25 | 6.624 | Yes |
| VvP45bXO-20 | 63 | 0.05 | 61.9 | 4.56 | 30 | 6 | 1.89 | Yes |
| VvP45bXO-21 | 133 | 0.05 | 74.4 | 9.34 | 74 | 3 | 9.443 | Yes |
| VvP45bXO-22 | 66 | 0.05 | 68.2 | 10.21 | 48 | 6 | 3.168 | Yes |
| VvP45bXO-23 | 66 | 0.05 | 68.2 | 6.7 | 42 | 6 | 2.772 | Yes |
| VvP45bXO-24 | 43 | 0.04 | 46.5 | 18.87 | 192 | 10 | 8.256 | Yes |
| VvP45bXO-25 | 63 | 0.04 | 64 | 7.53 | 36 | 6 | 2.268 | Yes |
| VvP45bXO-26 | 219 | 0.04 | 66.7 | 7.64 | 18 | 3 | 3.942 | Yes |

| | | | | | | | | |
|---|---|---|---|---|---|---|---|---|
| VvP45bXO-27 | 120 | 0.04 | 59.2 | 7.56 | 26 | 4 | 3.12 | Yes |
| VvP45bXO-28 | 128 | 0.04 | 59.4 | 4.78 | 27 | 3 | 3.456 | Yes |
| VvP45bXO-29 | 91 | 0.04 | 64.8 | 6.98 | 20 | 4 | 1.82 | Yes |
| VvP45bXO-30 | 149 | 0.03 | 56.4 | 13.54 | 58 | 3 | 8.642 | Yes |
| VvP45bXO-31 | 121 | 0.03 | 65.3 | 18.97 | 19 | 3 | 2.299 | Yes |
| VvP45bXO-32 | 52 | 0.03 | 46.2 | 8.44 | 77 | 5 | 8.008 | Yes |
| VvP45bXO-33 | 79 | 0.03 | 54.8 | 7.44 | 47 | 6 | 3.713 | Yes |
| VvP45bXO-34 | 1710 | 0.03 | 59.4 | 6 | 25 | 1 | 42.75 | Yes |
| VvP45bXO-35 | 26 | 0.03 | 61.5 | 17.43 | 212 | 16 | 5.512 | Yes |
| VvP45bXO-36 | 54 | 0.02 | 51.9 | 10.51 | 49 | 8 | 2.646 | Yes |
| VvP45bXO-37 | 139 | 0.02 | 62.2 | 5.34 | 15 | 3 | 2.445 | Yes |
| VvP45bXO-38 | 257 | 0.02 | 65 | 4.45 | 7 | 1 | 1.799 | Yes |
| VvP45bXO-39 | 271 | 0.02 | 69 | 12.6 | 24 | 2 | 1.704 | Yes |
| VvP45bXO-40 | 51 | 0.02 | 62.7 | 8.05 | 35 | 7 | 1.785 | Yes |
| VvP45bXO-41 | 68 | 0.02 | 55.9 | 3.92 | 25 | 5 | 1.7 | Yes |
| VvP45bXO-42 | 79 | 0.02 | 62 | 13.68 | 143 | 6 | 11.297 | Yes |
| VvP45bXO-43 | 110 | 0.02 | 47.3 | 5.84 | 48 | 4 | 5.28 | Yes |
| VvP45bXO-44 | 79 | 0.02 | 59.5 | 9.49 | 79 | 6 | 6.241 | Yes |
| VvP45bXO-45 | 1190 | 0.02 | 55.2 | 5.9 | 15 | 1 | 17.85 | Yes |
| VvP45bXO-46 | 129 | 0.02 | 53.5 | 11.98 | 124 | 5 | 15.996 | Yes |
| VvP45bXO-47 | 66 | 0.02 | 57.6 | 9.37 | 89 | 9 | 5.874 | Yes |
| VvP45bXO-48 | 22 | 0.02 | 63.6 | 14.22 | 1070 | 24 | 23.54 | Yes |
| VvP45bXO-49 | 52 | 0.02 | 47.1 | 10.86 | 179 | 9 | 9.308 | Yes |
| VvP45bXO-50 | 171 | 0.02 | 63.2 | 9.42 | 57 | 3 | 9.747 | Yes |
| VvP45bXO-51 | 1070 | 0.02 | 52.1 | 1.42 | 13 | 1 | 13.91 | Yes |
| VvP45bXO-52 | 13 | 0.01 | 46.2 | 9.44 | 570 | 30 | 7.41 | Yes |
| VvP45bXO-53 | 124 | 0.01 | 63.7 | 9.22 | 63 | 4 | 7.812 | Yes |
| VvP45bXO-54 | 61 | 0.01 | 65.6 | 10.01 | 211 | 9 | 12.871 | Yes |
| VvP45bXO-55 | 53 | 0.01 | 52.8 | 6.8 | 43 | 8 | 2.279 | Yes |
| VvP45bXO-56 | 108 | 0.01 | 61.1 | 4.57 | 23 | 4 | 2.484 | Yes |
| VvP45bXO-57 | 51 | 0.01 | 45.1 | 14.05 | 278 | 9 | 14.178 | Yes |
| VvP45bXO-58 | 86 | 0.01 | 52.3 | 4.94 | 24 | 4 | 2.064 | Yes |
| VvP45bXO-59 | 53 | 0.01 | 56.6 | 11.31 | 35 | 7 | 1.855 | Yes |
| VvP45bXO-60 | 32 | 0.01 | 40.6 | 10.24 | 122 | 12 | 3.904 | Yes |
| 5S rRNA + NTS | 151 | 0.01 | 49 | 11.87 | - | - | - | - |
| H2A histone | 376 | 0.0031 | 42.5 | 1.99 | - | - | - | - |

| | | | | | | | | |
|---|---|---|---|---|---|---|---|---|
| H4 histone | 378 | 0.0029 | 44.8 | 2.02 | - | - | - | - |
| H2B histone | 221 | 0.0017 | 43.8 | 2.66 | - | - | - | - |
| U2 snRNA | 199 | 0.0017 | 50.3 | 1.41 | - | - | - | - |
| H3 histone | 301 | 0.0016 | 42.7 | 1.5 | - | - | - | - |
| U1 snRNA | 162 | 0.00077 | 45.1 | 3.33 | - | - | - | - |
| U5 snRNA | 117 | 0.00052 | 58.1 | 5.33 | - | - | - | - |
| U6 snRNA | 107 | 0.00035 | 56.1 | 8.42 | - | - | - | - |

The satDNAs families and the telomere tandem repeat were assembled by RepeatExplorer2, and the tandem multigene families were assembled by NOVOPlasty. Abundance and divergence were estimated by RepeatMasker. satDNA families are named as VvP45XO; with "Vv" for *Vandiemenella viatica* group following by the "P" from provisional taxon name, and a number in a descending order of the genomic read proportion. A+T %: percentage of A+T in the consensus sequences. Kimura divergence %: Kimura 2-parameter divergence from the consensus sequence. Total of monomers in clusters: number of monomers recovered for each satDNA family in each the clusters in RepeatExplorer output. Max. number of monomers per tandem arrays per contig: maximum number of andemly repeated monomers found in the largest contig within the cluster. Total repeat family length (kb): length of each satDNA family in kb obtained by the total of monomers in clusters per consensus size. HORs: High order repeat structures identified by visual inspection in a dotplot.

**Table S5:** Most abundant tandem repeats (TRs) in the P45bXY sex chromosome race (male reads). The repeats are sorted in descending order of the genome proportion.

| Repeat name | Consensus length (bp) | Genome proportion % | A+T % | K2P divergence % | Total of monomers in clusters | Max. number of monomers per tandem arrays per contig | Total repeat family length (kb) | HORs |
|---|---|---|---|---|---|---|---|---|
| VvP45bXY-1 | 1250 | 1.20 | 55.1 | 23.67 | 39 | 1 | 48.75 | Yes |
| Telomere repeat | 5 | 1.19 | 60 | 3.45 | 499 | 65 | 2.495 | Yes |
| VvP45bXY-2 | 189 | 0.75 | 56 | 13.38 | 23 | 2 | 8.464 | Yes |
| VvP45bXY-3 | 167 | 0.44 | 66.5 | 18.3 | 317 | 4 | 52.939 | Yes |
| VvP45bXY-4 | 284 | 0.43 | 64.1 | 19.85 | 390 | 4 | 110.76 | Yes |
| VvP45bXY-5 | 101 | 0.39 | 49.5 | 8.31 | 78 | 4 | 7.878 | Yes |
| VvP45bXY-6 | 518 | 0.38 | 59.7 | 11.2 | 140 | 1 | 72.52 | Yes |
| 45S rRNA | 7876 | 0.32 | 43.2 | 1.43 | - | - | - | - |
| VvP45bXY-7 | 20 | 0.22 | 65 | 12.57 | 162 | 12 | 6.48 | Yes |
| VvP45bXY-8 | 31 | 0.21 | 54.8 | 7.42 | 114 | 12 | 3.534 | Yes |
| VvP45bXY-9 | 147 | 0.20 | 64.6 | 11.9 | 70 | 3 | 10.584 | Yes |
| VvP45bXY-10 | 101 | 0.20 | 56.4 | 8.73 | 65 | 4 | 7.767 | Yes |
| VvP45bXY-11 | 52 | 0.19 | 51.9 | 9.29 | 75 | 8 | 3.9 | Yes |
| VvP45bXY-12 | 99 | 0.16 | 60.6 | 8.21 | 51 | 6 | 5.049 | Yes |
| VvP45bXY-13 | 80 | 0.14 | 55 | 16.06 | 40 | 6 | 3.2 | Yes |
| VvP45bXY-14 | 22 | 0.12 | 40.9 | 8.42 | 288 | 19 | 6.336 | Yes |
| VvP45bXY-15 | 54 | 0.11 | 51.9 | 8.92 | 89 | 9 | 4.806 | Yes |
| VvP45bXY-16 | 16 | 0.10 | 50 | 10.29 | 777 | 28 | 12.432 | Yes |
| VvP45bXY-17 | 52 | 0.09 | 55.8 | 7.45 | 51 | 8 | 2.601 | Yes |
| VvP45bXY-18 | 32 | 0.09 | 65.6 | 4.38 | 84 | 12 | 2.688 | Yes |
| VvP45bXY-19 | 67 | 0.09 | 58.2 | 6.44 | 25 | 5 | 1.675 | Yes |
| VvP45bXY-20 | 34 | 0.08 | 61.8 | 4.36 | 70 | 10 | 2.38 | Yes |
| VvP45bXY-21 | 50 | 0.08 | 56 | 8.5 | 208 | 12 | 10.4 | Yes |
| VvP45bXY-22 | 34 | 0.08 | 61.8 | 4.01 | 114 | 16 | 3.876 | Yes |
| VvP45bXY-23 | 66 | 0.07 | 71.2 | 10.03 | 97 | 9 | 6.402 | Yes |
| VvP45bXY-24 | 66 | 0.06 | 65.2 | 11.26 | 42 | 6 | 2.772 | Yes |
| VvP45bXY-25 | 31 | 0.06 | 48.4 | 4.22 | 55 | 11 | 1.705 | Yes |
| VvP45bXY-26 | 51 | 0.06 | 51 | 19.86 | 319 | 12 | 16.269 | Yes |
| VvP45bXY-27 | 54 | 0.06 | 57.4 | 10.38 | 88 | 8 | 4.752 | Yes |
| VvP45bXY-28 | 91 | 0.06 | 64.8 | 7.34 | 20 | 4 | 1.82 | Yes |
| VvP45bXY-29 | 25 | 0.05 | 32 | 6.3 | 1300 | 23 | 32.5 | Yes |

| | | | | | | | | |
|---|---|---|---|---|---|---|---|---|
| VvP45bXY-30 | 66 | 0.05 | 56.1 | 8.31 | 40 | 10 | 2.64 | Yes |
| VvP45bXY-31 | 220 | 0.05 | 65 | 8.8 | 10 | 2 | 2.2 | Yes |
| VvP45bXY-32 | 52 | 0.05 | 46.2 | 6.82 | 99 | 8 | 5.148 | Yes |
| VvP45bXY-33 | 133 | 0.04 | 74.4 | 6.98 | 24 | 4 | 3.192 | Yes |
| VvP45bXY-34 | 122 | 0.04 | 63.1 | 12.36 | 38 | 3 | 4.636 | Yes |
| VvP45bXY-35 | 77 | 0.04 | 44.9 | 19.42 | 88 | 6 | 7.776 | Yes |
| VvP45bXY-36 | 75 | 0.04 | 45.3 | 7.78 | 55 | 5 | 4.125 | Yes |
| VvP45bXY-37 | 105 | 0.04 | 60 | 12.83 | 76 | 4 | 7.98 | Yes |
| VvP45bXY-38 | 35 | 0.03 | 60 | 5.05 | 82 | 12 | 2.87 | Yes |
| VvP45bXY-39 | 73 | 0.03 | 60.3 | 7.36 | 31 | 6 | 2.263 | Yes |
| VvP45bXY-40 | 62 | 0.03 | 67.7 | 14.27 | 130 | 9 | 8.06 | Yes |
| VvP45bXY-41 | 45 | 0.03 | 44.4 | 11.27 | 246 | 12 | 11.07 | Yes |
| VvP45bXY-42 | 64 | 0.03 | 57 | 11.3 | 148 | 9 | 9.472 | Yes |
| VvP45bXY-43 | 63 | 0.03 | 63.5 | 4.3 | 30 | 6 | 1.89 | Yes |
| VvP45bXY-44 | 63 | 0.03 | 55.6 | 8.05 | 66 | 6 | 4.158 | Yes |
| VvP45bXY-45 | 79 | 0.03 | 60.8 | 7.67 | 42 | 6 | 3.318 | Yes |
| VvP45bXY-46 | 16 | 0.03 | 37.5 | 11.72 | 493 | 25 | 7.888 | Yes |
| VvP45bXY-47 | 41 | 0.03 | 43.9 | 6.83 | 40 | 8 | 1.64 | Yes |
| VvP45bXY-48 | 50 | 0.02 | 52 | 10.18 | 146 | 9 | 7.3 | Yes |
| VvP45bXY-49 | 108 | 0.02 | 59.3 | 5.73 | 20 | 4 | 2.16 | Yes |
| VvP45bXY-50 | 19 | 0.02 | 42.1 | 11.26 | 957 | 27 | 18.183 | Yes |
| VvP45bXY-51 | 49 | 0.02 | 42.1 | 10.15 | 68 | 9 | 3.332 | Yes |
| VvP45bXY-52 | 66 | 0.02 | 56.1 | 7.41 | 63 | 9 | 4.158 | Yes |
| VvP45bXY-53 | 53 | 0.02 | 52.8 | 7.16 | 42 | 7 | 2.226 | Yes |
| VvP45bXY-54 | 20 | 0.02 | 35 | 9.82 | 184 | 18 | 3.68 | Yes |
| VvP45bXY-55 | 61 | 0.02 | 65.6 | 12.94 | 78 | 6 | 4.758 | Yes |
| VvP45bXY-56 | 47 | 0.02 | 57.4 | 5.46 | 35 | 7 | 1.645 | Yes |
| VvP45bXY-57 | 13 | 0.02 | 46.2 | 9.61 | 608 | 32 | 7.904 | Yes |
| VvP45bXY-58 | 189 | 0.02 | 51.3 | 9.78 | 15 | 3 | 2.835 | Yes |
| VvP45bXY-59 | 122 | 0.02 | 60.7 | 6.34 | 39 | 3 | 4.758 | Yes |
| VvP45bXY-60 | 64 | 0.02 | 62.5 | 7.64 | 36 | 6 | 2.304 | Yes |
| VvP45bXY-61 | 208 | 0.02 | 48.1 | 6.24 | 5 | 1 | 1.04 | Yes |
| VvP45bXY-62 | 121 | 0.02 | 59.5 | 5.92 | 35 | 4 | 4.235 | Yes |
| VvP45bXY-63 | 159 | 0.02 | 49.7 | 5.8 | 18 | 3 | 2.862 | Yes |
| VvP45bXY-64 | 34 | 0.01 | 50 | 7.8 | 94 | 12 | 3.196 | Yes |
| VvP45bXY-65 | 58 | 0.01 | 48.3 | 6.12 | 30 | 6 | 1.74 | Yes |
| VvP45bXY-66 | 1060 | 0.01 | 52 | 1.85 | 13 | 1 | 13.78 | Yes |
| VvP45bXY-67 | 50 | 0.01 | 56 | 4.58 | 35 | 7 | 1.75 | Yes |
| VvP45bXY-68 | 82 | 0.01 | 56.1 | 6.53 | 51 | 6 | 4.182 | Yes |

| | | | | | | | | |
|---|---|---|---|---|---|---|---|---|
| VvP45bXY-69 | 122 | 0.01 | 52.5 | 6.49 | 15 | 3 | 1.83 | Yes |
| VvP45bXY-70 | 45 | 0.01 | 35.6 | 6.72 | 177 | 10 | 7.965 | Yes |
| VvP45bXY-71 | 192 | 0.01 | 61.5 | 6.12 | 33 | 3 | 6.336 | Yes |
| VvP45bXY-72 | 183 | 0.01 | 60.1 | 4.33 | 24 | 3 | 4.392 | Yes |
| VvP45bXY-73 | 85 | 0.01 | 61.2 | 4.42 | 39 | 6 | 3.315 | Yes |
| VvP45bXY-74 | 181 | 0.01 | 45.3 | 8.14 | 24 | 3 | 4.344 | Yes |
| VvP45bXY-75 | 101 | 0.01 | 61.4 | 6.35 | 20 | 4 | 2.02 | Yes |
| 5S rRNA + NTS | 151 | 0.01 | 49.7 | 10.91 | - | - | - | - |
| VvP45bXY-76 | 110 | 0.01 | 41.8 | 7.95 | 24 | 4 | 2.64 | Yes |
| VvP45bXY-77 | 15 | 0.01 | 46.7 | 4.34 | 276 | 22 | 4.14 | Yes |
| VvP45bXY-78 | 1060 | 0.01 | 94.5 | 1.59 | 13 | 1 | 13.78 | Yes |
| VvP45bXY-79 | 123 | 0.01 | 48 | 5.94 | 40 | 4 | 4.92 | Yes |
| VvP45bXY-80 | 48 | 0.01 | 54.2 | 5.01 | 35 | 7 | 1.68 | Yes |
| VvP45bXY-81 | 24 | 0.01 | 41.7 | 3.62 | 188 | 26 | 4.512 | Yes |
| H3 histone | 411 | 0.003 | 40.4 | 1.52 | - | - | - | - |
| H2B histone | 372 | 0.003 | 45.2 | 1.92 | - | - | - | - |
| H2A histone | 338 | 0.0028 | 42.7 | 1.6 | - | - | - | - |
| U2 snRNA | 212 | 0.0017 | 51.9 | 1.18 | - | - | - | - |
| H4 histone | 111 | 0.0011 | 49.5 | 3.26 | - | - | - | - |
| U1 snRNA | 179 | 0.00046 | 49.2 | 3.52 | - | - | - | - |
| U6 snRNA | 108 | 0.00029 | 56.5 | 6.85 | - | - | - | - |
| U5 snRNA | 118 | 0.00008 | 55.1 | 5.62 | - | - | - | - |

The satDNA families and the telomere tandem repeat were assembled by RepeatExplorer2, and the tandem multigene families were assembled by NOVOPlasty. Abundance and divergence were estimated by RepeatMasker. satDNA families are named as VvP45XY; with "Vv" for *Vandiemenella viatica* group following by the "P" from provisional taxon name, and a number in a descending order of the genomic read proportion. A+T %: percentage of A+T in the consensus sequences. Kimura divergence %: Kimura 2-parameter from the consensus sequence. Total of monomers in clusters: number of monomers recovered for each satDNA family in each the clusters in RepeatExplorer output. Max. number of monomers per tandem arrays per contig: maximum number of tandemly repeated monomers found in the largest contig within the cluster. Total repeat family length (kb): length of each satDNA family in kb obtained by the total of monomers in clusters per consensus size. HORs: High order repeat structures identified by visual inspection in a dotplot.

**Table S6: Presence of 129 satDNA families across four chromosomal races of *viatica* species group.** Homology search based on 1) all-against-all RepeatMasker comparison of satDNA consensus sequences among races, and on 2) genomic read mapping against the consensus sequences as reference in RepeatMasker. Each column represents the library of satDNAs of the race, and each row indicates the common satDNAs between them. satDNAs highlighted in gray were used as references for read mapping. The signs positive (+) and negative (-) indicate presence or absence of the satDNA family in the race based on read mapping. Columns with satDNA name non highlighted in grey indicates homologous consensus sequences between races,  based on all-against-all RepeatMasker comparison of satDNA consensus sequences.

| RepeatMasker homology search | | | | |
|---|---|---|---|---|
| **Repeat** | **P24XO satDNA library** | **P24XY satDNA library** | **P45bXO satDNA library** | **P45bXY satDNA library** |
| satDNA-1 | VvP24XO-1 (51 bp) | VvP24XY-8 (51 bp) | VvP45bXO-7 (51 bp) | VvP45bXY-26 (51 bp) |
| satDNA-2 | VvP24XO-2 (53 bp) | + | VvP45bXO-59 (53 bp) | VvP45bXY-48 (50 bp) |
| satDNA-3 | VvP24XO-3 (257 bp) | + | VvP45bXO-38 (257 bp) | + |
| satDNA-4 | VvP24XO-5 (189 bp) | VvP24XY-1 (189 bp) | VvP45bXO-2 (189 bp) | VvP45bXY-2 (189 bp) |
| satDNA-5 | VvP24XO-6 (7 bp) | VvP24XY-7 (7bp) | + | + |
| satDNA-6 | VvP24XO-7 (147 bp) | VvP24XY-6 (147 bp) | VvP45bXO-9 (147 bp) | VvP45bXY-9 (147 bp) |
| satDNA-7 | VvP24XO-8 (66 bp) | - | + | VvP45bXY-30 (66 bp) |
| satDNA-8 | VvP24XO-9 (34 bp) | VvP24XY-42 (34 bp) | VvP45bXO-17 (34 bp) | VvP45bXY-22 (34 bp) |
| satDNA-9 | VvP24XO-10 (21 bp) | + | VvP45bXO-4 (21 bp) | + |
| satDNA-10 | VvP24XO-11 (125 bp) | VvP24XY-29 (131 bp) | VvP45bXO-28 (128 bp) | - |
| satDNA-11 | VvP24XO-12 (80 bp) | VvP24XY-9 (80 bp) | VvP45bXO-8 (80 bp) | VvP45bXY-13 (80 bp) |
| satDNA-12 | VvP24XO-13 (1,138 bp) | + | VvP45bXO-51 (1,066 bp) | VvP45bXY-66 (1,066 bp) |
| satDNA-13 | VvP24XO-14 (101 bp) | + | VvP45bXO-6 (101 bp) | VvP45bXY-10 (101 bp) |
| satDNA-14 | VvP24XO-15 (80 bp) | - | VvP45bXO-27 (80 bp) | - |
| satDNA-15 | VvP24XO-16 (75 bp) | VvP24XY-13 (75 bp) | + | VvP45bXY-36 (75 bp) |
| satDNA-16 | VvP24XO-17 (130 bp) | VvP24XY-10 (130 bp) | VvP45bXO-16 (130 bp) | + |
| satDNA-17 | VvP24XO-18 (51 bp) | + | VvP45bXO-3 (51 bp) | VvP24XY-2 (52 bp) |
| satDNA-18 | VvP24XO-19 (31 bp) | VvP24XY-16 (31 bp) | + | VvP45bXY-8 (31 bp) |

| | | | | |
|---|---|---|---|---|
| satDNA-19 | VvP24XO-20 (133 bp) | VvP24XY-11 (133 bp) | VvP45bXO-21 (133 bp) | VvP45bXY-33 (133 bp) |
| satDNA-20 | VvP24XO-21 (52 bp) | VvP24XY-15 (52 bp) | VvP45bXO-15 (52 bp) | VvP45bXY-17 (52 bp) |
| satDNA-21 | VvP24XO-22 (54 bp) | VvP24XY-38 (54 bp) | VvP45bXO-36 (54 bp) | VvP45bXY-27 (54 bp) |
| satDNA-22 | VvP24XO-23 (66 bp) | - | + | + |
| satDNA-23 | VvP24XO-24 (32 bp) | VvP24XY-24 (32 bp) | VvP45bXO-18 (32 bp) | VvP45bXY-18 (32 bp) |
| satDNA-24 | VvP24XO-25 (52 bp) | VvP24XY-17 (52 bp) | VvP45bXO-49 (52 bp) | VvP45bXY-32 (52 bp) |
| satDNA-25 | VvP24XO-26 (220 bp) | VvP24XY-18 (220 bp) | VvP45bXO-26 ((221 bp) | VvP45bXY-31 (221 bp) |
| satDNA-26 | VvP24XO-27 (79 bp) | VvP24XY-27 (79 bp) | VvP45bXO-33 (79 bp) | VvP45bXY-45 (79 bp) |
| satDNA-27 | VvP24XO-28 (52 bp) | VvP24XY-20 (53 bp) | VvP45bXO-32 (52 bp) | VvP45bXY-11 (52 bp) |
| satDNA-28 | VvP24XO-29 (26 bp) | VvP24XY-23 (26 bp) | + | + |
| satDNA-29 | VvP24XO-30 (131 bp) | VvP24XY-14 (130 bp) | VvP45bXO-16 (130bp) | + |
| satDNA-30 | VvP24XO-31 (63 bp) | VvP24XY-34 (63 bp) | VvP45bXO-20 (63 bp) | VvP45bXY-43 (63 bp) |
| satDNA-31 | VvP24XO-32 (122 bp) | + | VvP45bXO-31 (121 bp) | VvP45bXY-34 (122 bp) |
| satDNA-32 | VvP24XO-33 (160 bp) | VvP24XY-30 (160 bp) | VvP45bXO-50 (171) | + |
| satDNA-33 | VvP24XO-34 (122 bp) | VvP24XY-32 (123 bp) | VvP45bXO-53 (123 bp) | VvP45bXY-59 (122 bp) |
| satDNA-34 | VvP24XO-35 (42 bp) | VvP24XY-45 42 bp) | - | - |
| satDNA-35 | VvP24XO-36 (60 bp) | VvP24XY-37 (60 bp) | - | + |
| satDNA-36 | VvP24XO-37 (108 bp) | VvP24XY-44 (108 bp) | VvP45bXO-56 (108 bp) | VvP45bXY-49 (108 bp) |
| satDNA-37 | VvP24XO-38 (83 bp) | - | - | - |
| satDNA-38 | VvP24XO-39 (138 bp) | VvP24XY-33 (138 bp) | + | + |
| satDNA-39 | VvP24XO-40 (86 bp) | + | VvP45bXO-58 (86 bp) | + |
| satDNA-40 | VvP24XO-41 (26 bp) | + | + | + |
| satDNA-41 | VvP24XO-42 (33 bp) | + | + | + |
| satDNA-42 | VvP24XO-43 (76 bp) | + | + | + |
| satDNA-43 | VvP24XO-44 (36 bp) | VvP24XY-39 (36 bp) | - | - |

| | | | | |
|---|---|---|---|---|
| satDNA-44 | VvP24XO-45 (65 bp) | + | + | VvP45bXY-60 (64 bp) |
| satDNA-45 | + | VvP24XY-3 | + | + |
| satDNA-46 | + | VvP24XY-4 (168 bp) | + | VvP45bXY-3 (168 bp) |
| satDNA-47 | + | VvP24XY-5 (19 bp) | + | + |
| satDNA-48 | + | VvP24XY-12 (43 bp) | VvP45bXO-24 (43 bp) | + |
| satDNA-49 | + | VvP24XY-19 930 bp) | + | + |
| satDNA-50 | + | VvP24XY-21 (43 bp) | + | + |
| satDNA-51 | + | VvP24XY-22 (61 bp) | VvP45bXO-54 (61 bp) | VvP45bXY-40 (62 bp) |
| satDNA-52 | + | VvP24XY-25 (1,069 bp) | + | + |
| satDNA-53 | + | VvP24XY-26 (68 bp) | VvP45bXO-41 (68 bp) | + |
| satDNA-54 | + | VvP24XY-28 (66 bp) | VvP45bXO-47 (66 bp) | VvP45bXY-52 (66 bp) |
| satDNA-55 | + | VvP24XY-31 (1,690 bp) | VvP45bXO-34 (1,710 bp) | + |
| satDNA-56 | + | VvP24XY-35 (20 bp) | VvP45bXO-11 (20 bp) | VvP45bXY-7 (20 bp) |
| satDNA-57 | - | VvP24XY-36 (121 bp) | + | VvP45bXY-62 (121 bp) |
| satDNA-58 | - | VvP24XY-40 (104 bp) | - | - |
| satDNA-59 | - | VvP24XY-41 (29 bp) | - | - |
| satDNA-60 | + | VvP24XY-43 (32 bp) | + | + |
| satDNA-61 | - | VvP24XY-46 (77 bp) | + | VvP45bXY-77 (77 bp) |
| satDNA-62 | + | VvP24XY-47 (19 bp) | + | + |
| satDNA-63 | + | VvP24XY-48 (42 bp) | - | - |
| satDNA-64 | + | + | VvP45bXO-1 | + |
| satDNA-65 | + | + | VvP45bXO-5 (282 bp) | VvP45bXY-4 (284 bp) |
| satDNA-66 | - | + | VvP45bXO-10 (54 bp) | VvP45bXY-15 (54 bp) |
| satDNA-67 | - | - | VvP45bXO-12 (63 bp) | - |
| satDNA-68 | + | + | VvP45bXO-13 (99 bp) | VvP45bXY-12 (99 bp) |

| | | | | |
|---|---|---|---|---|
| satDNA-69 | + | + | VvP45bXO-14 (67 bp) | VvP45bXY-19 (67 bp) |
| satDNA-70 | + | + | VvP45bXO-19 (16 bp) | VvP45bXY-16 (16 bp) |
| satDNA-71 | + | + | VvP45bXO-22 (66 bp) | VvP45bXY-23 (66 bp) |
| satDNA-72 | - | + | VvP45bXO-23 (66 bp) | VvP45bXY-24 (66 bp) |
| satDNA-73 | - | + | VvP45bXO-25 (63 bp) | VvP45bXY-44 (63 bp) |
| satDNA-74 | + | + | VvP45bXO-29 (91 bp) | VvP45bXY-28 (91 bp) |
| satDNA-75 | + | + | VvP45bXO-30 (149 bp) | + |
| satDNA-76 | + | + | VvP45bXO-35 (26 bp) | + |
| satDNA-77 | - | - | VvP45bXO-37 | + |
| satDNA-78 | - | - | VvP45bXO-39 | - |
| satDNA-79 | + | + | VvP45bXO-40 | + |
| satDNA-80 | + | + | VvP45bXO-42 | - |
| satDNA-81 | + | + | VvP45bXO-43 | + |
| satDNA-82 | + | + | VvP45bXO-44 | + |
| satDNA-83 | + | + | VvP45bXO-45 (1,190 bp) | + |
| satDNA-84 | + | - | VvP45bXO-46 | - |
| satDNA-85 | + | + | VvP45bXO-48 | + |
| satDNA-86 | + | + | VvP45bXO-52 (13 bp) | VvP45bXY-57(13 bp) |
| satDNA-87 | + | + | VvP45bXO-55 (53 bp) | VvP45bXY-53 (53 bp) |
| satDNA-88 | + | + | VvP45bXO-57 (51 bp) | + |
| satDNA-89 | + | + | VvP45bXO-60 | + |
| satDNA-90 | + | + | + | VvP45bXY-1 (1,250 bp) |
| satDNA-91 | + | + | + | VvP45bXY-5 (101 bp) |
| satDNA-92 | + | + | + | VvP45bXY-6 (518 bp) |
| satDNA-93 | + | + | + | VvP45bXY-14 (22 bp) |

| | | | | |
|---|---|---|---|---|
| satDNA-94 | + | + | + | VvP45bXY-20 (34 bp) |
| satDNA-95 | + | + | + | VvP45bXY-21 (50 bp) |
| satDNA-96 | - | - | + | VvP45bXY-25 (31 bp) |
| satDNA-97 | + | + | + | VvP45bXY-29 (25 bp) |
| satDNA-98 | + | + | + | VvP45bXY-35 (77 bp) |
| satDNA-99 | + | + | + | VvP45bXY-37 (105 bp) |
| satDNA-100 | + | + | + | VvP45bXY-38 (35 bp) |
| satDNA-101 | - | - | - | VvP45bXY-39 (73 bp) |
| satDNA-102 | + | + | + | VvP45bXY-41 (45 bp) |
| satDNA-103 | - | - | + | VvP45bXY-42 (64 bp) |
| satDNA-104 | + | + | + | VvP45bXY-46 (16 bp) |
| satDNA-105 | + | + | + | VvP45bXY-47 (41 bp) |
| satDNA-106 | + | + | + | VvP45bXY-50 (19 bp) |
| satDNA-107 | + | + | + | VvP45bXY-51 (49 bp) |
| satDNA-108 | + | + | + | VvP45bXY-54 (20 bp) |
| satDNA-109 | + | + | + | VvP45bXY-55 (61 bp) |
| satDNA-110 | + | + | + | VvP45bXY-56 (47 bp) |
| satDNA-111 | + | + | + | VvP45bXY-58 (189 bp) |
| satDNA-112 | + | + | + | VvP45bXY-61 (208 bp) |
| satDNA-113 | + | + | + | VvP45bXY-63 (159 bp) |
| satDNA-114 | + | - | + | VvP45bXY-64 (34 bp) |
| satDNA-115 | + | + | + | VvP45bXY-65 (58 bp) |
| satDNA-116 | + | + | + | VvP45bXY-67 (50 bp) |
| satDNA-117 | + | + | + | VvP45bXY-68 (82 bp) |
| satDNA-118 | + | + | + | VvP45bXY-69 (122 bp) |

| | | | | |
|---|---|---|---|---|
| satDNA-119 | + | + | + | VvP45bXY-70 (45 bp) |
| satDNA-120 | + | + | + | VvP45bXY-71 (192 bp) |
| satDNA-121 | + | + | + | VvP45bXY-72 (183 bp) |
| satDNA-122 | + | + | + | VvP45bXY-73 (85 bp) |
| satDNA-123 | + | + | + | VvP45bXY-74 (181 bp) |
| satDNA124 | - | + | - | VvP45bXY-75 (101) |
| satDNA-125 | + | + | + | VvP45bXY-76 (110 bp) |
| satDNA-126 | + | + | + | VvP45bXY-78 (1,060) |
| satDNA-127 | + | - | + | VvP45bXY-79 (123 bp) |
| satDNA-128 | + | + | + | VvP45bXY-80 (48 bp) |
| satDNA-129 | + | + | + | VvP45bXY-81 (24 bp) |

**Table S7:** The number of satDNA families that were present in only one of the races or absent in at least one race.

| Present only in | Total satDNA families (repeat names) |
|---|---|
| P24X0 | 1 (satDNA-37) |
| P24XY | 2 (satDNA-58, satDNA-59) |
| P45bX0 | 2 (satDNA-67, satDNA-78) |
| P45bXY | 1 (satDNA-101) |
| **Absent in** | |
| P24X0 | 5 (satDNA-57, satDNA-61, satDNA-66, satDNA-72, satDNA-73) |
| P24XY | 4 (satDNA-7, satDNA-22, satDAN-114, satDNA-127) |
| P45bX0 | 1 (satDNA-35) |
| P45bXY | 2 (satDAN-10, satDNA-80) |
| P24 races | 3 (satDNA-77, satDNA-96-satDNA-103) |
| P45b races | 3 (satDNA-34, satDNA-43, satDNA-63) |
| X0 races | 1 (satDNA-124) |
| XY races | 2 (satDNA-14, satDNA-84) |

**Table S8: Statistics of the 102 satDNA families shared across the four chromosomal races of the *viatica* species group. 50 satDNA families show differential amplification in copy number (cv > 80 %). GP = Genome proportion. K2P = Kimura 2-parameter distance. sd = standard deviation. cv = coefficient of variation.**

| Family | P24X0 | | P24XY | | P45bX0 | | P45bXY | | GP stats | | | K2P stats | |
|---|---|---|---|---|---|---|---|---|---|---|---|---|---|
| | GP % | K2P % | GP % | K2P % | GP % | K2P % | GP % | K2P % | Average | sd | cv % | Average | sd |
| satDNA-1 | 1.48 | 13.94 | 0.13 | 19.6 | 0.23 | 20.28 | 0.06 | 19.86 | 0.47 | 0.67 | 141.8 | 18.42 | 2.99 |
| satDNA-2 | 0.96 | 7.12 | 0.33 | 9.5 | 0.01 | 11.31 | 0.02 | 10.18 | 0.33 | 0.44 | 135.0 | 9.53 | 1.77 |
| satDNA-3 | 0.94 | 5.24 | 0.0002 | 4.54 | 0.02 | 4.45 | 0.008 | 13.22 | 0.24 | 0.46 | 192.26 | 6.86 | 4.25 |
| satDNA-4 | 0.89 | 6.3 | 0.88 | 11.96 | 0.77 | 12.79 | 0.75 | 13.38 | 0.82 | 0.07 | 8.84 | 11.10 | 3.25 |
| satDNA-5 | 0.74 | 21.1 | 0.19 | 13.65 | 0.4 | 19.63 | 0.49 | 18.52 | 0.45 | 0.22 | 50.07 | 18.22 | 3.22 |
| satDNA-6 | 0.32 | 14.02 | 0.26 | 18.77 | 0.18 | 13.79 | 0.2 | 11.9 | 0.24 | 0.06 | 26.35 | 14.62 | 2.92 |
| satDNA-8 | 0.17 | 8.11 | 0.01 | 4.72 | 0.06 | 6.57 | 0.08 | 4.01 | 0.08 | 0.06 | 83.54 | 5.85 | 1.85 |
| satDNA-9 | 0.15 | 5.1 | 0.03 | 12.48 | 0.46 | 8.05 | 0.7 | 12.38 | 0.33 | 0.30 | 90.56 | 9.5 | 3.58 |
| satDNA-11 | 0.07 | 5.31 | 0.08 | 7.23 | 0.21 | 8.31 | 0.14 | 16.06 | 0.12 | 0.06 | 51.64 | 9.23 | 4.72 |
| satDNA-12 | 0.07 | 6.5 | 0.07 | 4.66 | 0.02 | 1.42 | 0.01 | 1.85 | 0.04 | 0.03 | 75.33 | 3.61 | 2.40 |
| satDNA-13 | 0.07 | 3.16 | 0.31 | 23.93 | 0.38 | 8.53 | 0.19 | 8.905 | 0.24 | 0.13 | 57.46 | 11.13 | 8.92 |
| satDNA-15 | 0.07 | 10.1 | 0.05 | 7.35 | 0.005 | 19.04 | 0.04 | 7.78 | 0.04 | 0.02 | 65.93 | 11.07 | 5.45 |
| satDNA-16 | 0.07 | 7.27 | 0.05 | 6.58 | 0.06 | 5.96 | 9E-05 | 34.46 | 0.04 | 0.03 | 68.96 | 13.56 | 13.94 |
| satDNA-17 | 0.06 | 5.4 | 0.86 | 26.75 | 0.5 | 15.77 | 0.35 | 11.98 | 0.44 | 0.33 | 75.23 | 14.97 | 8.94 |
| satDNA-18 | 0.06 | 24.4 | 0.04 | 6.85 | 0.0002 | 18.88 | 0.21 | 7.42 | 0.08 | 0.09 | 118.29 | 14.39 | 8.67 |
| satDNA-19 | 0.06 | 6.91 | 0.05 | 5.23 | 0.05 | 9.34 | 0.04 | 6.98 | 0.05 | 0.008 | 16.33 | 7.11 | 1.68 |
| satDNA-20 | 0.05 | 8.22 | 0.04 | 7.36 | 0.07 | 7.07 | 0.09 | 7.45 | 0.06 | 0.02 | 35.48 | 7.52 | 0.49 |
| satDNA-21 | 0.04 | 6.46 | 0.04 | 9.1 | 0.02 | 10.51 | 0.06 | 10.38 | 0.04 | 0.02 | 40.82 | 9.11 | 1.88 |
| satDNA-23 | 0.04 | 7.9 | 0.02 | 7.1 | 0.06 | 6.73 | 0.09 | 4.38 | 0.05 | 0.03 | 56.88 | 6.53 | 1.51 |
| satDNA-24 | 0.03 | 6.67 | 0.04 | 5.66 | 0.02 | 10.86 | 0.05 | 6.82 | 0.03 | 0.01 | 36.89 | 7.50 | 2.29 |
| satDNA-25 | 0.03 | 8.01 | 0.04 | 7.72 | 0.04 | 7.64 | 0.05 | 8.8 | 0.04 | 0.008 | 20.41 | 8.04 | 0.53 |
| satDNA-26 | 0.03 | 10.71 | 0.02 | 7.31 | 0.03 | 7.44 | 0.03 | 7.67 | 0.03 | 0.005 | 18.18 | 8.28 | 1.62 |
| satDNA-27 | 0.03 | 7.43 | 0.03 | 9.51 | 0.03 | 8.44 | 0.19 | 9.29 | 0.07 | 0.08 | 114.29 | 8.67 | 0.94 |
| satDNA-28 | 0.02 | 6.73 | 0.02 | 5.46 | 0.03 | 15.77 | 0.02 | 18.87 | 0.02 | 0.005 | 22.22 | 11.7 | 6.62 |
| satDNA-29 | 0.02 | 11.73 | 0.05 | 5.41 | 0.06 | 5.96 | 0.04 | 5.7 | 0.04 | 0.02 | 40.18 | 7.2 | 3.03 |
| satDNA-30 | 0.02 | 7.82 | 0.01 | 4.27 | 0.05 | 4.56 | 0.03 | 4.3 | 0.03 | 0.02 | 62.10 | 5.24 | 1.73 |
| satDNA-31 | 0.02 | 7.94 | 0.02 | 11.62 | 0.03 | 18.97 | 0.04 | 12.36 | 0.03 | 0.009 | 34.82 | 12.72 | 4.59 |

| | | | | | | | | | | | | | |
|---|---|---|---|---|---|---|---|---|---|---|---|---|---|
| satDNA-32 | 0.02 | 13.66 | 0.02 | 6.45 | 0.02 | 9.42 | 0.02 | 8 | 0.02 | 0 | 0.00 | 9.38 | 3.09 |
| satDNA-33 | 0.02 | 8.36 | 0.02 | 5.74 | 0.01 | 9.22 | 0.02 | 6.34 | 0.02 | 0.005 | 28.57 | 7.41 | 1.64 |
| satDNA-36 | 0.01 | 16.72 | 0.01 | 3.15 | 0.01 | 4.57 | 0.02 | 5.73 | 0.01 | 0.005 | 40.00 | 7.54 | 6.21 |
| satDNA-38 | 0.01 | 3.36 | 0.02 | 3.69 | 0.015 | 2.35 | 0.01 | 2.35 | 0.01 | 0.005 | 34.82 | 2.94 | 0.69 |
| satDNA-39 | 0.01 | 6.7 | 0.007 | 8.47 | 0.01 | 4.94 | 0.005 | 9.15 | 0.008 | 0.002 | 30.62 | 7.31 | 1.89 |
| satDNA-40 | 0.01 | 4.02 | 0.005 | 10.77 | 0.006 | 7.97 | 0.0006 | 11.22 | 0.005 | 0.004 | 71.50 | 8.49 | 3.31 |
| satDNA-41 | 0.01 | 6.91 | 0.02 | 26.3 | 0.01 | 27.69 | 0.05 | 28.54 | 0.02 | 0.02 | 84.13 | 22.36 | 10.34 |
| satDNA-42 | 0.01 | 10.17 | 0.003 | 7.19 | 0.004 | 8.31 | 7E-05 | 26.96 | 0.004 | 0.004 | 92.29 | 13.16 | 9.28 |
| satDNA-44 | 0.01 | 11.84 | 0.015 | 11 | 0.015 | 7.62 | 0.02 | 7.64 | 0.015 | 0.004 | 27.22 | 9.52 | 2.21 |
| satDNA-45 | 0.01 | 3.68 | 0.53 | 17.38 | 0.32 | 23.58 | 0.51 | 25.53 | 0.34 | 0.24 | 70.37 | 17.54 | 9.87 |
| satDNA-46 | 0.51 | 16.65 | 0.51 | 16.42 | 0.365 | 17.12 | 0.44 | 18.3 | 0.45 | 0.07 | 15.17 | 17.12 | 0.83 |
| satDNA-47 | 0.0002 | 11.57 | 0.27 | 10.01 | 0.03 | 9.83 | 0.005 | 11.14 | 0.08 | 0.13 | 170.11 | 10.64 | 0.85 |
| satDNA-48 | 0.01 | 20.1 | 0.05 | 16.18 | 0.04 | 18.87 | 0.06 | 18.34 | 0.04 | 0.022 | 54.01 | 18.37 | 1.64 |
| satDNA-49 | 5E-05 | 8.22 | 0.03 | 5.58 | 5E-05 | 10.12 | 3E-05 | 9.37 | 0.007 | 0.01 | 198.85 | 8.32 | 1.98 |
| satDNA-50 | 0.005 | 18.8 | 0.03 | 17.17 | 0.04 | 19.45 | 0.05 | 19.45 | 0.03 | 0.02 | 61.80 | 18.72 | 1.07 |
| satDNA-51 | 0.013 | 10.01 | 0.03 | 8.47 | 0.01 | 10.01 | 0.013 | 7.12 | 0.016 | 0.009 | 54.16 | 8.9 | 1.39 |
| satDNA-52 | 0.02 | 13.79 | 0.02 | 7.92 | 0.015 | 17.39 | 0.018 | 14.94 | 0.02 | 0.002 | 12.95 | 13.51 | 4.01 |
| satDNA-53 | 7E-05 | 0.52 | 0.02 | 5.46 | 0.007 | 1.87 | 0.007 | 1.5 | 0.008 | 0.008 | 98.85 | 2.34 | 2.15 |
| satDNA-54 | 0.004 | 25.13 | 0.02 | 11.24 | 0.02 | 9.37 | 0.02 | 7.41 | 0.016 | 0.008 | 50.00 | 13.28 | 8.04 |
| satDNA-55 | 0.01 | 11.16 | 0.02 | 4.84 | 0.03 | 6.48 | 0.02 | 11.63 | 0.02 | 0.008 | 40.82 | 8.53 | 3.38 |
| satDNA-56 | 0.004 | 11.33 | 0.01 | 10.71 | 0.13 | 12.89 | 0.22 | 12.57 | 0.09 | 0.10 | 114.01 | 11.87 | 1.02 |
| satDNA-60 | 0.007 | 24.18 | 0.01 | 22.67 | 0.05 | 30.01 | 0.08 | 30.33 | 0.04 | 0.03 | 94.87 | 26.79 | 3.94 |
| satDNA-62 | 1E-05 | 5.71 | 0.01 | 7.33 | 0.0003 | 3.73 | 0.001 | 6.48 | 0.003 | 0.005 | 169.75 | 5.81 | 1.54 |
| satDNA-64 | 0.64 | 21.25 | 0.8 | 14.63 | 0.89 | 21.68 | 0.88 | 21.76 | 0.8 | 0.11 | 14.39 | 19.83 | 3.47 |
| satDNA-65 | 0.48 | 22.3 | 0.41 | 21.94 | 0.44 | 19.87 | 0.43 | 19.85 | 0.44 | 0.03 | 6.69 | 20.99 | 1.31 |
| satDNA-68 | 0.01 | 18.17 | 0.001 | 13.16 | 0.09 | 10.25 | 0.16 | 8.21 | 0.06 | 0.07 | 114.59 | 12.45 | 4.32 |
| satDNA-69 | 0.006 | 8.26 | 0.006 | 9.43 | 0.08 | 6.16 | 0.09 | 6.44 | 0.04 | 0.04 | 100.64 | 7.57 | 1.55 |
| satDNA-70 | 0.007 | 27.76 | 0.01 | 28.16 | 0.06 | 8.49 | 0.1 | 10.29 | 0.04 | 0.04 | 100.36 | 18.67 | 10.75 |
| satDNA-71 | 5E-05 | 29.81 | 4E-05 | 32.03 | 0.05 | 10.21 | 0.07 | 10.03 | 0.03 | 0.03 | 118.46 | 20.52 | 12.04 |
| satDNA-74 | 0.006 | 14.08 | 0.001 | 13.5 | 0.04 | 6.98 | 0.06 | 7.34 | 0.03 | 0.03 | 105.18 | 10.47 | 3.84 |
| satDNA-75 | 0.02 | 16.72 | 0.0175 | 18.23 | 0.03 | 13.54 | 0.02 | 18.39 | 0.02 | 0.005 | 25.34 | 16.72 | 2.24 |

| | | | | | | | | | | | | | |
|---|---|---|---|---|---|---|---|---|---|---|---|---|---|
| satDNA-76 | 0.02 | 14.5 | 0.03 | 16.39 | 0.03 | 17.43 | 0.01 | 24.19 | 0.02 | 0.009 | 42.55 | 18.13 | 4.21 |
| satDNA-79 | 0.006 | 16.46 | 0.006 | 14.37 | 0.02 | 8.05 | 0.008 | 15.01 | 0.01 | 0.006 | 67.33 | 13.47 | 3.72 |
| satDNA-81 | 0.002 | 7.62 | 0.002 | 6.06 | 0.02 | 13.68 | 0.006 | 6.82 | 0.007 | 0.008 | 113.92 | 8.54 | 3.48 |
| satDNA-82 | 8E-05 | 20.59 | 7E-05 | 26.48 | 0.02 | 5.84 | 8E-05 | 21.42 | 0.005 | 0.009 | 196.97 | 18.58 | 8.88 |
| satDNA-83 | 0.008 | 7.28 | 0.008 | 8.56 | 0.02 | 9.49 | 0.01 | 7 | 0.01 | 0.006 | 49.95 | 8.08 | 1.16 |
| satDNA-85 | 0.04 | 13.91 | 0.02 | 14.29 | 0.02 | 11.98 | 0.009 | 16.11 | 0.02 | 0.012 | 58.07 | 14.07 | 1.69 |
| satDNA-86 | 0.003 | 14.66 | 0.003 | 13.82 | 0.02 | 14.22 | 0.02 | 9.61 | 0.01 | 0.009 | 85.35 | 13.08 | 2.34 |
| satDNA-87 | 0.0002 | 33.44 | 0.008 | 6.51 | 0.01 | 9.44 | 0.02 | 7.16 | 0.009 | 0.008 | 85.33 | 14.14 | 12.93 |
| satDNA-88 | 0.006 | 14.95 | 0.001 | 17.74 | 0.01 | 6.8 | 0.001 | 16.43 | 0.004 | 0.004 | 96.86 | 13.98 | 4.92 |
| satDNA-89 | 0.0004 | 18.26 | 0.0003 | 16.77 | 0.01 | 14.05 | 0.004 | 12.3 | 0.004 | 0.004 | 123.93 | 15.34 | 2.67 |
| satDNA-90 | 1 | 22.82 | 1 | 23.02 | 1.2 | 23.67 | 1.2 | 23.67 | 1.1 | 0.11 | 10.50 | 23.29 | 0.44 |
| satDNA-91 | 1.14 | 27.27 | 1.2 | 20.61 | 0.73 | 15.88 | 0.39 | 8.31 | 0.9 | 0.37 | 43.85 | 18.02 | 7.98 |
| satDNA-92 | 0.44 | 17.04 | 0.4 | 16.57 | 0.46 | 17.26 | 0.38 | 11.2 | 0.42 | 0.04 | 8.69 | 15.51 | 2.89 |
| satDNA-93 | 0.003 | 19.24 | 0.003 | 20.03 | 0.006 | 14.98 | 0.12 | 8.42 | 0.033 | 0.06 | 175.81 | 15.67 | 5.31 |
| satDNA-94 | 0.14 | 26.13 | 0.01 | 25.18 | 0.06 | 25.71 | 0.08 | 4.36 | 0.07 | 0.05 | 74.17 | 20.34 | 10.67 |
| satDNA-95 | 9E-05 | 23.66 | 0.0006 | 23.75 | 7E-05 | 20.32 | 0.08 | 8.5 | 0.02 | 0.04 | 197.49 | 19.05 | 7.21 |
| satDNA-97 | 0.002 | 13.1 | 0.0007 | 12.01 | 0.008 | 10.38 | 0.05 | 6.3 | 0.01 | 0.02 | 154.42 | 10.45 | 2.98 |
| satDNA-98 | 0.03 | 24.36 | 0.02 | 22.57 | 0.03 | 23.6 | 0.04 | 19.42 | 0.03 | 0.008 | 27.22 | 22.49 | 2.17 |
| satDNA-99 | 0.005 | 19.03 | 0.01 | 19.12 | 0.07 | 22.13 | 0.04 | 12.83 | 0.03 | 0.03 | 96.33 | 18.28 | 3.91 |
| satDNA-100 | 0.0009 | 8.75 | 0.0003 | 7.49 | 0.003 | 6.35 | 0.03 | 5.05 | 0.008 | 0.01 | 167.80 | 6.91 | 1.58 |
| satDNA-102 | 0.002 | 10.79 | 0.001 | 11.47 | 0.001 | 11.83 | 0.03 | 14.27 | 0.008 | 0.01 | 168.72 | 12.09 | 1.51 |
| satDNA-104 | 0.0004 | 18 | 0.0007 | 18.12 | 0.003 | 17.31 | 0.03 | 11.3 | 0.008 | 0.01 | 168.49 | 16.18 | 3.27 |
| satDNA-105 | 0.003 | 19.76 | 0.002 | 16.64 | 0.003 | 5.28 | 0.03 | 11.72 | 0.009 | 0.01 | 143.95 | 13.35 | 6.31 |
| satDNA-106 | 0.006 | 10.79 | 0.006 | 11.74 | 0.008 | 11.2 | 0.03 | 6.83 | 0.01 | 0.01 | 93.64 | 10.14 | 2.24 |
| satDNA-107 | 0.001 | 17.59 | 0.0005 | 13.07 | 0.001 | 17.59 | 0.02 | 11.26 | 0.006 | 0.009 | 170.42 | 14.88 | 3.22 |
| satDNA-108 | 0.001 | 15.99 | 0.0007 | 23.5 | 0.0006 | 16.47 | 0.02 | 10.15 | 0.005 | 0.009 | 172.52 | 16.53 | 5.46 |
| satDNA-109 | 0.001 | 24.81 | 0.03 | 19.36 | 0.01 | 24.58 | 0.02 | 9.82 | 0.01 | 0.01 | 82.14 | 19.64 | 7.01 |
| satDNA-110 | 0.0004 | 14.19 | 0.0003 | 10.64 | 0.0004 | 14.5 | 0.02 | 12.94 | 0.005 | 0.009 | 186.10 | 13.07 | 1.75 |
| satDNA-111 | 0.004 | 22.46 | 0.007 | 15.34 | 0.005 | 22.84 | 0.02 | 5.46 | 0.009 | 0.007 | 82.65 | 16.52 | 8.14 |
| satDNA-112 | 0.003 | 15.09 | 0.002 | 12.98 | 0.003 | 15.17 | 0.02 | 9.78 | 0.007 | 0.008 | 123.99 | 13.25 | 2.523 |
| satDNA-113 | 0.002 | 27.33 | 0.001 | 19.46 | 0.002 | 18.76 | 0.02 | 6.24 | 0.006 | 0.009 | 146.86 | 17.95 | 8.72 |

| | | | | | | | | | | | | | |
|---|---|---|---|---|---|---|---|---|---|---|---|---|---|
| satDNA-115 | 0.002 | 10.67 | 0.005 | 9.84 | 0.001 | 17 | 0.01 | 7.8 | 0.004 | 0.004 | 89.81 | 11.33 | 3.96 |
| satDNA-116 | 0.004 | 4.38 | 0.0004 | 4.7 | 0.0001 | 5.84 | 0.01 | 6.12 | 0.004 | 0.005 | 127.02 | 5.26 | 0.85 |
| satDNA-117 | 0.004 | 6.35 | 0.006 | 4.63 | 0.006 | 5.66 | 0.01 | 4.58 | 0.006 | 0.002 | 38.72 | 5.30 | 0.85 |
| satDNA-118 | 0.006 | 10.46 | 0.002 | 10.8 | 0.001 | 11.73 | 0.01 | 6.53 | 0.005 | 0.004 | 86.59 | 9.88 | 2.29 |
| satDNA-119 | 8E-05 | 21.45 | 0.0005 | 20.97 | 0.0005 | 14.35 | 0.01 | 6.49 | 0.003 | 0.005 | 174.15 | 15.81 | 7.01 |
| satDNA-120 | 0.006 | 27.41 | 0.005 | 26.23 | 0.008 | 24.26 | 0.01 | 6.72 | 0.007 | 0.002 | 30.58 | 21.15 | 9.71 |
| satDNA-121 | 0.0004 | 5.81 | 0.0002 | 7.59 | 0.002 | 3.9 | 0.01 | 6.12 | 0.003 | 0.005 | 147.21 | 5.855 | 1.52 |
| satDNA-122 | 0.03 | 40.05 | 0.0001 | 25.71 | 0.0003 | 34.05 | 0.01 | 4.33 | 0.01 | 0.014 | 139.09 | 26.03 | 15.62 |
| satDNA-123 | 0.0003 | 30.46 | 0.0004 | 30.12 | 0.006 | 11.73 | 0.01 | 4.42 | 0.004 | 0.005 | 112.79 | 19.18 | 13.17 |
| satDNA-125 | 0.004 | 9.91 | 0.0007 | 13.43 | 0.006 | 11.63 | 0.01 | 6.35 | 0.005 | 0.004 | 75.15 | 10.33 | 3.02 |
| satDNA-126 | 0.008 | 3 | 0.004 | 10.01 | 0.003 | 3.36 | 0.01 | 7.95 | 0.006 | 0.003 | 52.86 | 6.08 | 3.45 |
| satDNA-128 | 2E-05 | 10.27 | 0.009 | 2.69 | 0.006 | 5.8 | 0.01 | 5.94 | 0.006 | 0.004 | 71.79 | 6.17 | 3.11 |
| satDNA-129 | 0.003 | 17.85 | 0.001 | 10.11 | 0.0004 | 11.79 | 0.01 | 5.01 | 0.003 | 0.004 | 122.47 | 11.19 | 5.29 |
