## Additional file 2 for "Too much too many: comparative analysis of morabine grasshopper genomes reveals highly abundant transposable elements and rapidly proliferating satellite DNA repeats"

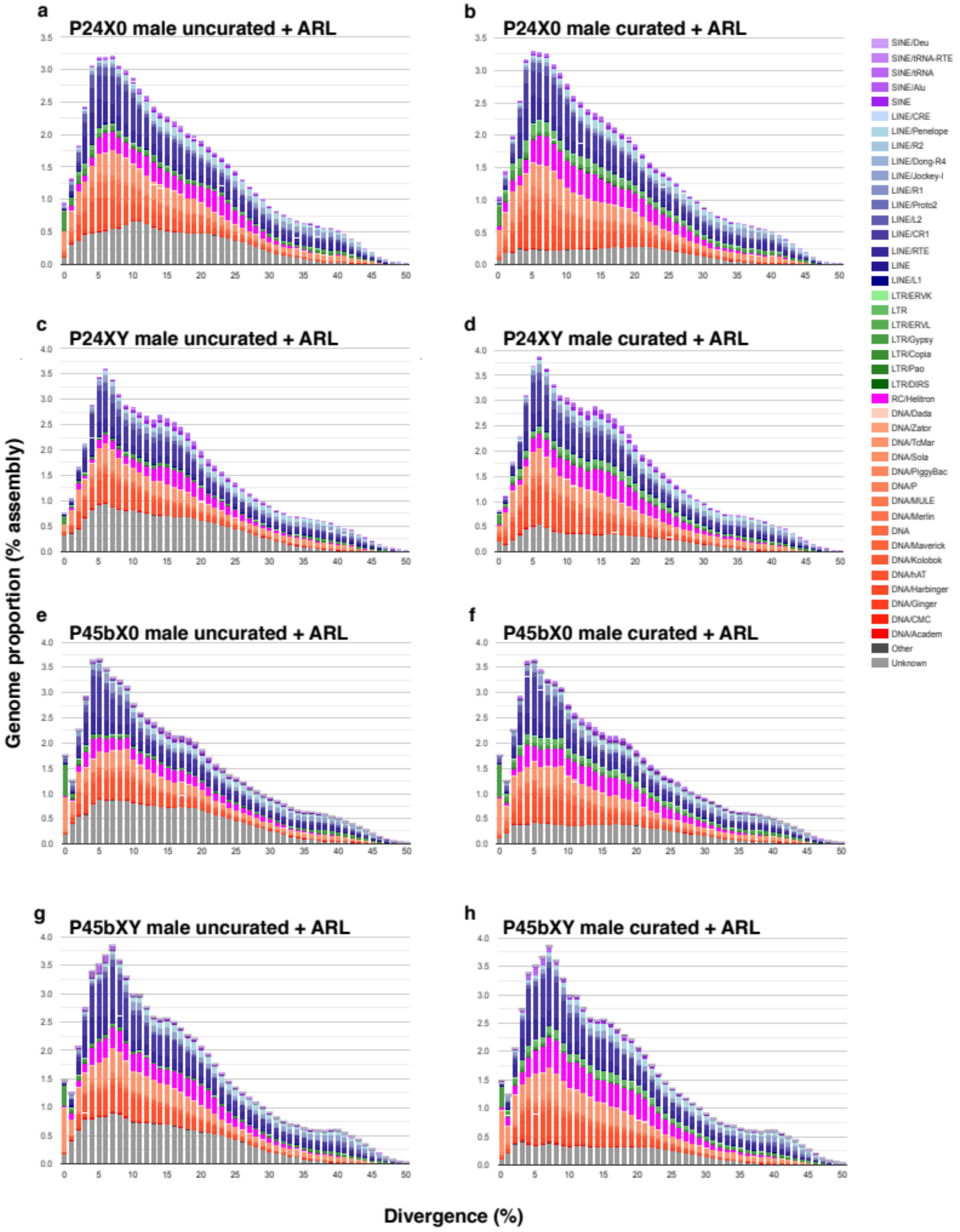


**Figure S1: Comparison of transposable element landscapes in male genome assemblies of four chromosomal races of the *viatica* species group. a-h)** Percentage of bp occupied in the genome (*y* axis) plotted against the Kimura 2-parameter (transitions/transversions) distance (*x* axis) of copies from each TE superfamily (color-coded) from their consensus sequences. **a,c,e,g)** Based on *de-novo* predicted repeats from RepeatModeler (RML) and Arthropoda Repbase library (*A*RL) repeats. **b,d,f,h)** Based on curated *de-novo* predicted repeats (curation of the P24X0 library used for re-classification of the three other *de-novo* libraries) from RepeatModeler + *A*RL repeats. Note the share of unknown (grey) repeats, a majority of which were identified as LTR retrotransposons (green) and DNA transposons (orange/red) when manually curated.

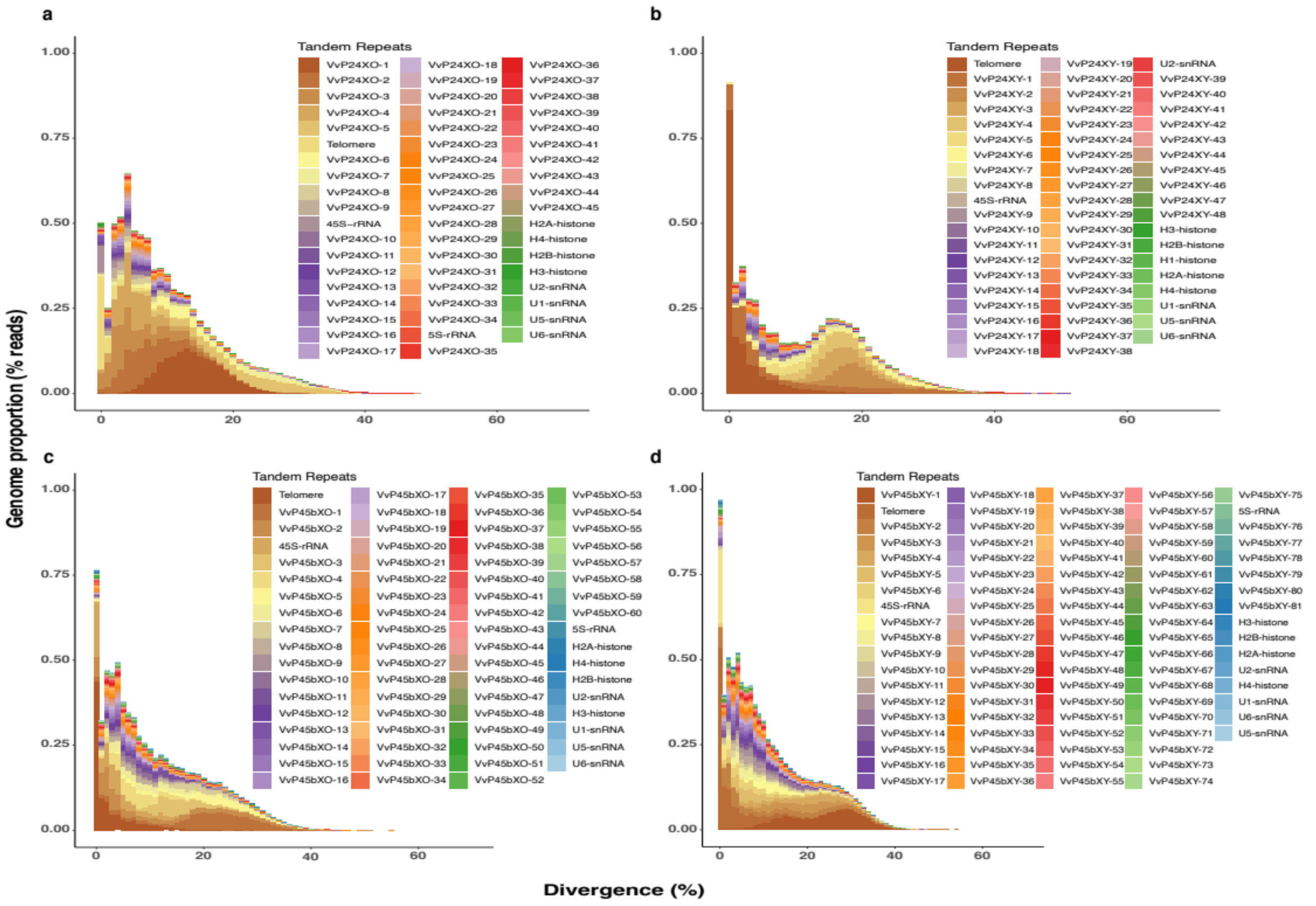


**Figure Figure S2: Tandem repeat (TR) landscapes in the male sequenced reads of four chromosomal races of the *viatica* species group. a)** P24XO. **b)** P24XY. **c)** P45bXO. **d)** P45bXY. Temporal accumulation of TRs is shown as repeat element divergence in Kimura-2 parameter (K2P) distance to consensus on the *x* axis and the TR abundance in on the *y* axis. The satDNAs are named as "Vv" (for *Vandiemenella viatica* group) followed by "P" (for provisional taxon) and a number that indicates the family number in decreasing order of the genomic read proportion of the race.

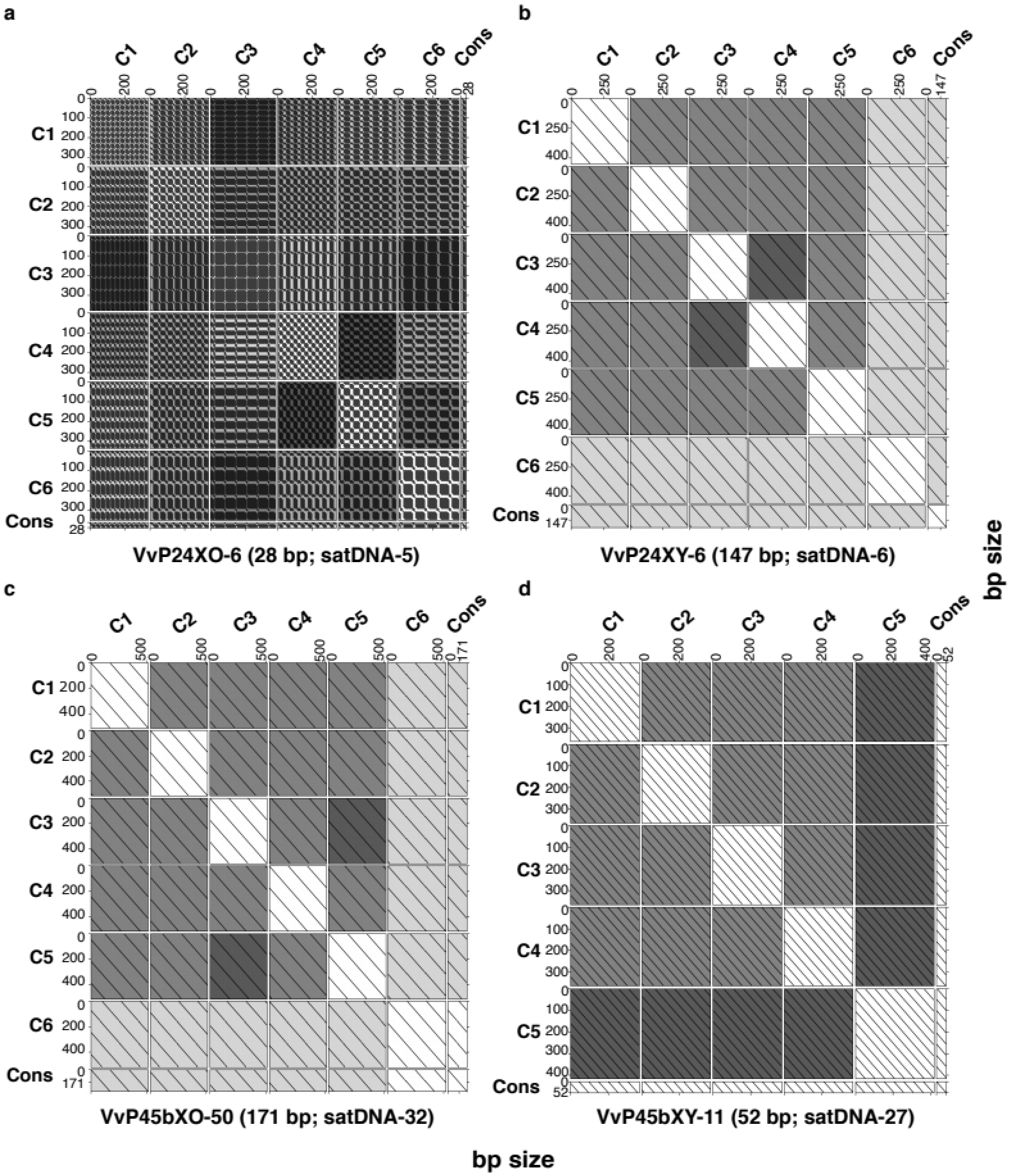


***Figure S3: All-against-all dotplot comparisons showing the diversity of satDNAs arrays detected among four chromosomal races of the viatica* species group. a)** The VvP24XO-6 (28 bp) satDNA in the P24XO race. **b)** The VvP24XY-6 (167 bp) satDNA in the P24XY race. **c)** The VvP45bXO-50 (171 bp) satDNA in the P45bXO race. **d)** The VvP45bXY-11 (52 bp) satDNA in the P45bXY race. The satDNAs are named as "Vv" (for *Vandiemenella viatica* group) followed by "P" (for provisional taxon) and a number that indicates the family number in decreasing order of the genomic read proportion of the race. Each satDNA family was defined by graph-based clustering of sequencing reads in RepeatExplorer2 (see Materials and Methods section). For simplicity, the plot shows all-against-all comparisons of the first six contigs (C1-C6) within the cluster. Contigs are aligned against themselves, against one another, and against their monomer consensus sequence (Cons). The different grey/black shades enable the identification of long shared subsequences between contigs at a glance, based on longest common subsequence, or longest match if mismatches are considered. Longer matches are represented by darker background shading.
